# Fetal microglia show region-specific and morphology-dependent sex differences in their responsiveness to prenatal maternal stress

**DOI:** 10.64898/2026.08.14.744921

**Authors:** Alexandra Lawson, Matthew Rosin, Jessica M. Rosin

## Abstract

The prevalence of neurodevelopmental disorders (NDDs) has increased dramatically, with growing evidence linking prenatal maternal stress exposure to NDDs. Across diverse maternal stressors, immune dysregulation emerges as a common feature, suggesting that fetal microglia may detect changes in the intrauterine environment and influence neurodevelopment. Accordingly, we utilized a mouse model of prenatal maternal cold stress to investigate the impact of maternal stress during pregnancy on fetal microglia morphology, cellular interactions, and phagocytic behaviors. Pregnant mice were exposed to cold stress from embryonic day 11.5 (E11.5) to E15.5 and fetal hypothalamic tissue was assessed from both male and female embryos. By adapting the morphology analysis toolset MicrogliaMorphology to assess fetal microglia, we demonstrate regional differences in microglial morphology in the fetal hypothalamus at baseline, with hypothalamic nuclei such as the paraventricular nucleus (PVN) containing fewer rod-like microglia compared to the broader hypothalamus. Interestingly, prenatal maternal cold stress induced a male-specific shift in microglial morphology from ameboid to ramified within the E15.5 PVN. Male embryos also displayed increased microglial-arginine vasopressin (AVP) neuronal interactions and microglial phagocytosis within the E15.5 PVN, but these changes were unique to microglia with a ramified morphology and were not observed when microglia with an ameboid or rod-like morphology were assessed. Using pHrodo bioparticles and flow cytometry, we further illustrate that prenatal maternal cold stress drives increased phagocytic activity in the E15.5 hypothalamus of male embryos, but not females. Together, these data demonstrate that prenatal maternal cold stress alters microglia morphology and drives morphology-dependent microglial interactions and phagocytic behaviors in male embryos which are unique to the hypothalamic PVN—a nuclei critical for social behaviors. Our findings also suggest that specific hypothalamic nuclei such as the PVN may be more sensitive to prenatal maternal stress, which has the potential to provide a cellular basis underlying the sex differences in microglia-dependent social deficits that were previously reported for this model.

## Background

During embryogenesis, the brain is highly plastic and the developmental trajectories of neural cells are particularly sensitive to cues from the intrauterine environment (1–5). A wide range of prenatal maternal stressors can disrupt this environment, including adverse psychological states, physiological challenges such as infection and inflammation, and environmental exposures (1,6–16). Despite their diverse origins, prenatal maternal stressors are consistently associated with altered fetal brain development and increased risk of neurodevelopmental disorders (NDDs), including autism spectrum disorder (ASD), attention-deficit/hyperactivity disorder (ADHD), anxiety, depression, and schizophrenia (1,8,13–15,17–22). While NDDs frequently show sex differences in their onset, presentation, and/or prevalence (23–31), the sex-divergent mechanisms that may underly these differences are poorly understood.

Microglia, which derive from primitive myeloid progenitors in the embryonic yolk sac rather than from central nervous system (CNS) progenitors, colonize the brain beginning around embryonic day 9.5 (E9.5) (32–37). This places microglia within the fetal brain during critical neurogenic and gliogenic programs, where they have been shown to act as key mediators of both pre- and postnatal neurodevelopment by actively shaping the early neural environment through regulation of neuronal and glial differentiation, axonal outgrowth, synaptic pruning, and clearance of apoptotic cells (1,38–43). As microglia express glucocorticoid receptors and are highly responsive to inflammatory environments (38,44–49), their presence in the fetal brain at E9.5 makes mid-gestation onwards a sensitive window where altered maternal stress signaling and immune dysregulation have the potential to cause enduring effects on the developing fetal brain (32,34,40,50–52). Although overt sex differences in microglial numbers and morphology have yet to be documented in the fetal brain and are suggested to arise postnatally (53–57), microglial responsiveness to intrauterine disruptions during pregnancy have been shown to differ between sexes (1,9,12,58–60).

As the hypothalamus acts as a bridge between the nervous and endocrine systems, it is not surprising that it is particularly tuned to sense alterations in stress signaling. Among the many diverse nuclei the comprise the hypothalamus, the paraventricular nucleus (PVN) is particularly relevant as it is a central integrator of stress and neuroendocrine responses and a source of oxytocin (OXT) and arginine vasopressin (AVP), peptides that have been directly implicated in ASD (61–64). Structurally, the hypothalamus itself is sexually dimorphic, and it is known to drive sex differences in reproductive and endocrine functions, in addition to behavior (65–69). Intriguingly, neurodevelopmental disruptions that arise in response to perturbation of the intrauterine environment, which often differ by sex and show lasting behavioral consequences in offspring, are common in the fetal hypothalamus (1,49,70–76), potentially positioning it as a brain region through which sex differences in NDDs may arise.

Among a variety of prenatal maternal stressors that have been utilized in animal models, prenatal maternal cold stress has been shown to elevate maternal corticosterone, induce placental inflammation, and alter fetal microglial physiology (1,11,77,78). Unlike more severe prenatal maternal stressors that can cause embryo reabsorption, exposure to prenatal maternal cold stress does not significantly affect litter size (1,11). Most relevant, recent work in the E15.5 hypothalamus has identified a population of fetal microglia that are located near and directly in contact with neural stem and progenitor cells (NSPCs) that are transcriptionally responsive to prenatal maternal cold stress. Moreover, prenatal maternal cold stress drove microglia-dependent changes in neural numbers and behavior, including a significant reduction in OXT neuron numbers in the E15.5 PVN and social deficits, but only in males (1). While female offspring exposed to prenatal maternal cold stress also developed social deficits, this effect was independent of microglia and appears to arise as a result of disruptions to NSPCs (76), suggesting that the cellular underpinnings underlying the social deficits observed in this prenatal maternal stress model may differ between males and females.

Despite these findings, transcriptional signatures only capture one dimension of microglial state, and it remains unclear how prenatal maternal cold stress exposure influences fetal hypothalamic microglial morphology, cellular interactions, and phagocytic behaviors. By adapting the MicrogliaMorphology toolset generated by Kim et al. (2024) to assess fetal microglia (79), we conduct spatially resolved morphological clustering to assess microglial morphology across the male and female hypothalamus, along the entire rostro-caudal axis, and within two specific nuclei, the PVN and arcuate nucleus (ARC), both at baseline and in response to prenatal maternal cold stress exposure. Moreover, we link these morphologic phenotypes to functional readouts by assessing microglial interactions with AVP-neurons and microglial phagocytosis within the PVN. Lastly, we utilize pHrodo bioparticles and flow cytometry to evaluate the phagocytic capacity of microglia in the E15.5 hypothalamus of both sexes, under baseline conditions and in response to prenatal maternal cold stress. Our findings show that microglial morphology displays regional differences in the fetal hypothalamus at baseline, and that prenatal maternal cold stress exposure alters microglial morphology, interactions with AVP neurons, and phagocytosis uniquely in males and females, providing insight into the cellular mechanisms underlying the effects of early-life maternal stress during pregnancy, and further, providing a cellular basis for how these diverse mechanisms could contribute to the sex differences observed across many NDDs.

## Results

### Microglial morphology displays regional differences in the fetal hypothalamus

During embryogenesis, microglia undergo transitions in morphology, shifting from an ameboid state towards increasingly ramified phenotypes as the brain matures (80,81); however, microglial morphology also varies across brain regions in the developing brain (53). It is also noteworthy that brain regions such as the hypothalamus are highly heterogeneous structures, composed of distinct nuclei with specialized functions, and as such, may exhibit region-specific differences in microglial morphology (34). Interestingly, prenatal maternal cold stress exposure has been shown to drive both sex-specific and microglia-dependent disruptions in the embryonic hypothalamus (1). While these findings highlight sex differences in microglial responsiveness to maternal stress in the embryonic brain, the specific cellular mechanisms involved in these changes remain unclear. As microglial morphology can serve as a proxy for functional state, it represents a valuable approach to begin understanding microglial cellular dynamics, particularly when combined with functional readouts (82). Accordingly, we utilized our previously established prenatal maternal cold stress model, whereby pregnant dams are subjected to 30 minutes of 4°C cold exposure daily from E11.5 to E15.5 (Figure 1A). Serial sectioning was performed in the coronal plane across the rostro-caudal axis of E15.5 control and cold stress male and female brains in order to assess microglial morphology in all regions of the hypothalamus, including the PVN and ARC as prenatal maternal cold stress exposure has been shown to alter offspring weight gain and social behaviors (1). Thresholding of each section in Fiji/ImageJ revealed thousands of microglia, with a total of 35,089 unique microglial cells analyzed across the hypothalamus of E15.5 control and cold stress brains, of which 16,970 were derived from male embryos and 18,119 from female embryos (Figure 1B). Interestingly, ∼5% of these hypothalamic microglia localized within the PVN, with a total of 1,928 unique microglial cells analyzed, of which 873 were derived from male embryos and 1,055 from female embryos (Figure 1C). In contrast, ∼14% of hypothalamic microglia localized to the ARC, with a total of 5,025 unique microglial cells analyzed, of which 2,427 were derived from male embryos and 2,598 from female embryos (Figure 1D).

**Figure 1.**
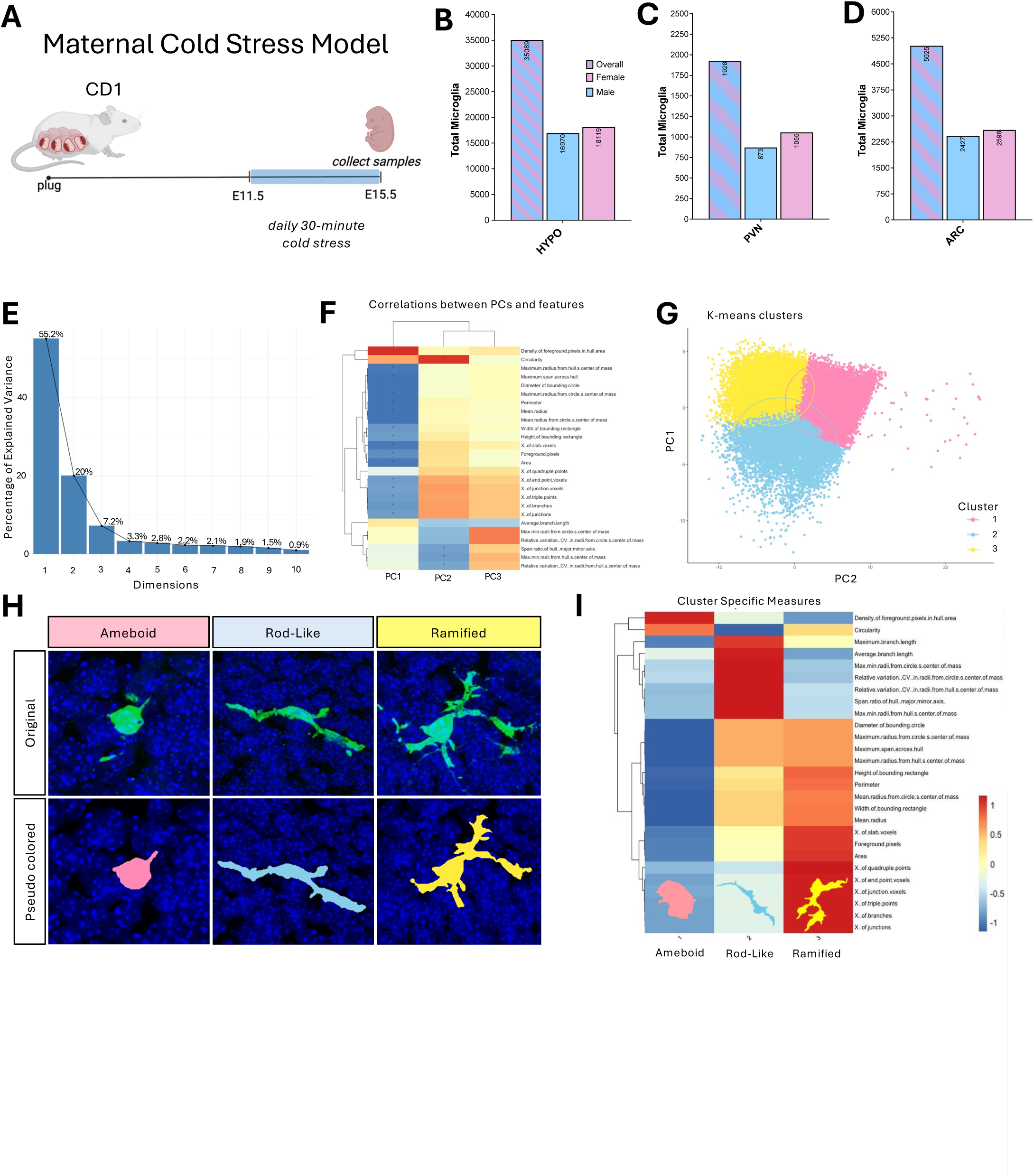
Experimental design and clustering of microglial cells. (A) Schematic illustrating the prenatal maternal cold stress paradigm. (B–D) Total number of microglia collected from the E15.5 hypothalamus (HYPO) (B), paraventricular nucleus (PVN) (C), and arcuate nucleus (ARC) (D). (E) Elbow plot showing the proportion of variance explained by the first 10 principal components. (F) Heatmap of pairwise Pearson correlations among extracted microglial morphological features. (G) Principal component analysis (PCA) for all microglia extracted. (H) Representative images of individual microglia from each identified cluster (ameboid, ramified, and rod-like) and their corresponding pseudo–colored reconstructions. (I) Heatmap summarizing the relationship between the 27 individual morphological features extracted for each microglial cell and cluster identity. Microglia were extracted from n=6 embryos per sex/treatment from 3 independent litters.

To characterize individual microglial morphology, single cells were extracted, skeletonized, and quantified by adapting the MicrogliaMorphology pipeline published by Kim et al. (2024). This analysis yielded 27 unique morphological features per cell, including features such as branch points, cell area, and cell circularity. Given that a single 2D morphological feature often fails to fully capture the unique complexity of microglial morphology (83), principal component analysis (PCA) was applied to reduce the 27 extracted features to lower-dimensional PC space. As over 75% of the variance was explained by the first two PCs, these were used for all downstream analyses (Figure 1E, F). K-means clustering was subsequently applied (Figure 1G), revealing three distinct morphological clusters: ameboid, rod-like, and ramified (Figure 1H). The ameboid cluster was characterized by high circularity and foreground pixel density within the hull area, with fewer branches and a smaller perimeter (Figure 1I). The rod-like cluster was distinguished by low circularity, a long average branch length, and few branch points (Figure 1I). The ramified cluster was marked by a high number of branch points, a short average branch length, and a large perimeter and cell area (Figure 1I).

Since the hypothalamus is a highly heterogeneous structure, and as such, likely exhibits region-specific differences in microglial morphology, we began by characterizing microglial morphology under control conditions. Cluster prevalence in the PVN, ARC, and whole hypothalamus was compared in E15.5 control male and female embryos combined, where microglia were pseudo-coloured by cluster identity (Figures 2A–C). A significant effect of cluster was observed (Figure 2D; cluster, χ²(2)=196.86, p<0.0001), consistent with the three morphological phenotypes being present in unequal proportions overall, with ramified and ameboid clusters comprising a greater proportion of microglia than the rod-like cluster. As this effect reflects the mathematical structure of k-means clustering, it appeared consistently across all cluster proportion-based analyses. Accordingly, we focused our analyses on within-cluster regional comparisons to determine whether the prevalence of each morphological cluster differed across hypothalamic regions. Microglial morphological cluster proportions varied significantly across hypothalamic regions in E15.5 control embryos (Figure 2D; cluster×brain region interaction, χ²(4)=20.98, p=0.0003). This difference appeared to be driven by the rod-like microglial cluster, as the whole hypothalamus was found to contain a significantly higher proportion of rod-like microglia relative to the PVN (Figure 2D; p=0.0029) and also showed a trend toward an increase in comparison to the ARC (Figure 2D; p=0.0873). In contrast, no difference was observed in the prevalence of the rod-like microglial cluster between the PVN and ARC (Figure 2D). Interestingly, ameboid and ramified microglial clusters showed no differences across each of the hypothalamic regions assessed, with no significant differences observed in ameboid cluster proportions or ramified cluster proportions (Figure 2D).

**Figure 2.**
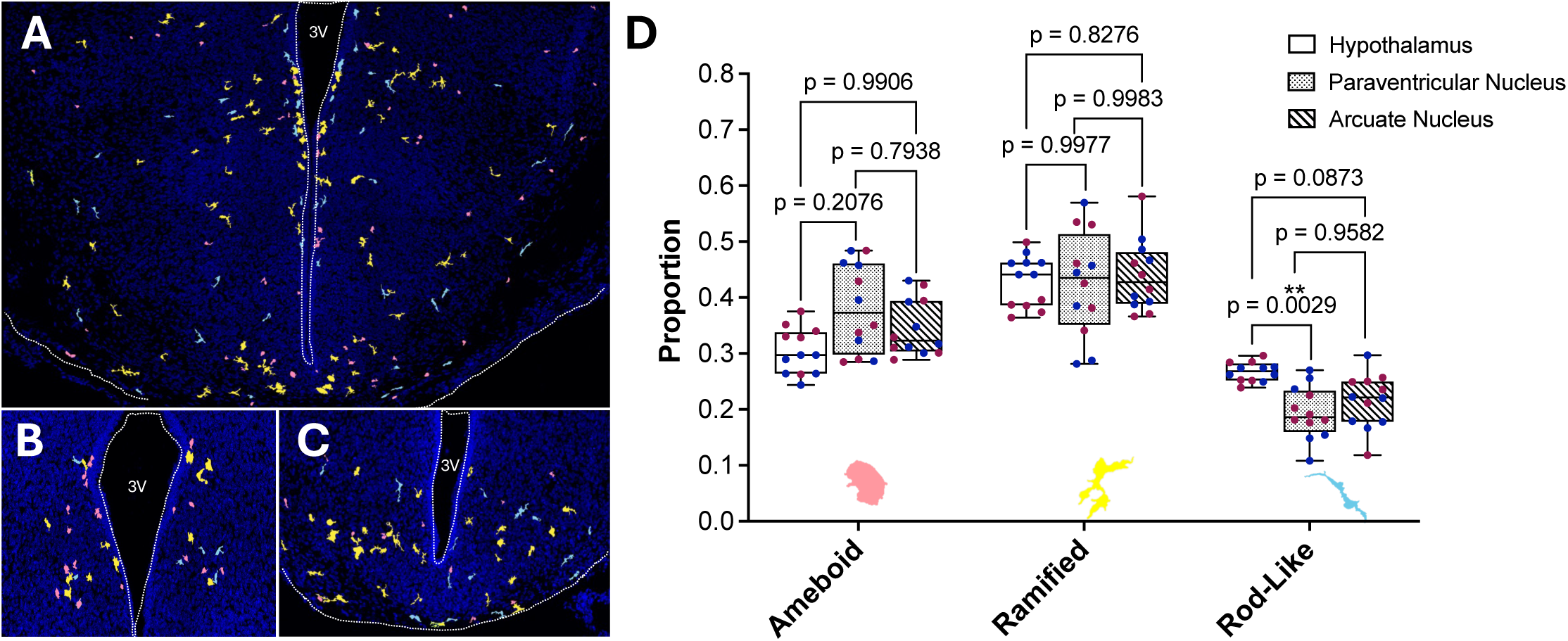
Microglial cluster prevalence in E15.5 embryos across the PVN, ARC, and whole hypothalamus. (A–C) Representative images of E15.5 control hypothalamic (A), PVN (B), and ARC (C) sections with ameboid (pink), ramified (yellow), and rod-like (blue) microglia pseudo–colored by cluster. (D) Proportion of each microglial cluster (ameboid vs. ramified vs. rod-like) across the E15.5 control male and female hypothalamus, PVN, and ARC (cluster, p*<*0.0001; brain region, p*=*0.9845; cluster×brain region interaction, p*=*0.0003). The proportion of rod-like microglia is significantly higher in the hypothalamus relative to the PVN (p*=*0.0029). Data are represented as box-and-whisker plots showing the median (center line), interquartile range (box), and minimum–maximum values (whiskers) and were analyzed by a linear mixed-effects model with Sidak-adjusted post hoc comparisons. Each dot represents an individual embryo (n=12), which were collected from 3 independent litters. Color indicates sex (blue, male; pink, female).

Together, these findings highlight region-specific differences in microglial cluster distribution within the developing hypothalamus, prompting for a region-specific approach for subsequent assessment of microglial morphology in control and prenatal maternal cold stress embryos.

### Prenatal maternal cold stress exposure drives sex- and region-specific changes in microglial morphology in the fetal hypothalamus

We next wanted to assess microglial morphology in the E15.5 hypothalamus, stratified by section across the rostro–caudal axis, and in the PVN and ARC, both at baseline and in response to prenatal maternal cold stress exposure. Again, we focused our analyses on within-cluster comparisons to determine whether the prevalence of each microglial morphological state differed across groups. To start, we examined microglia in E15.5 control and prenatal maternal cold stress male and female embryos across the rostro-caudal axis of the hypothalamus (Figure 3A–D), where we observed that the total number of microglia in the hypothalamus did not differ by sex or in response to prenatal maternal cold stress exposure (Figure 3E, Table S1). To examine if sex differences in microglial morphology exist, we first compared microglial cluster prevalence in the E15.5 male and female hypothalamus under control conditions. Consistent with previous analysis, we observed a significant effect of cluster; however, there was no impact of sex on microglial cluster proportions in the hypothalamus under control conditions (Figure 3F; cluster, χ²(2)=49.7396, p<0.0001). To better assess potential regional differences across the rostro–caudal axis of the E15.5 male and female hypothalamus under control conditions, each of the 12 sections spanning the entirety of the hypothalamus were examined separately. In section 1, microglial cluster proportions appeared to differ in a sex-dependent manner (Figure S1A, Table S2; cluster×sex interaction, χ²(2)=6.445, p=0.0399); however, this did not resolve to any specific cluster, with ameboid, ramified, and rod-like microglial cluster proportions showing no differences between control male and female embryos (Figure S1A). While a significant effect of cluster was observed for all hypothalamic sections (Table S2; cluster, χ²(2)=13.35, p=0.0013), no other hypothalamic section showed significant baseline sex differences in microglial cluster proportions (Figure S1A; Table S2).

**Figure 3.**
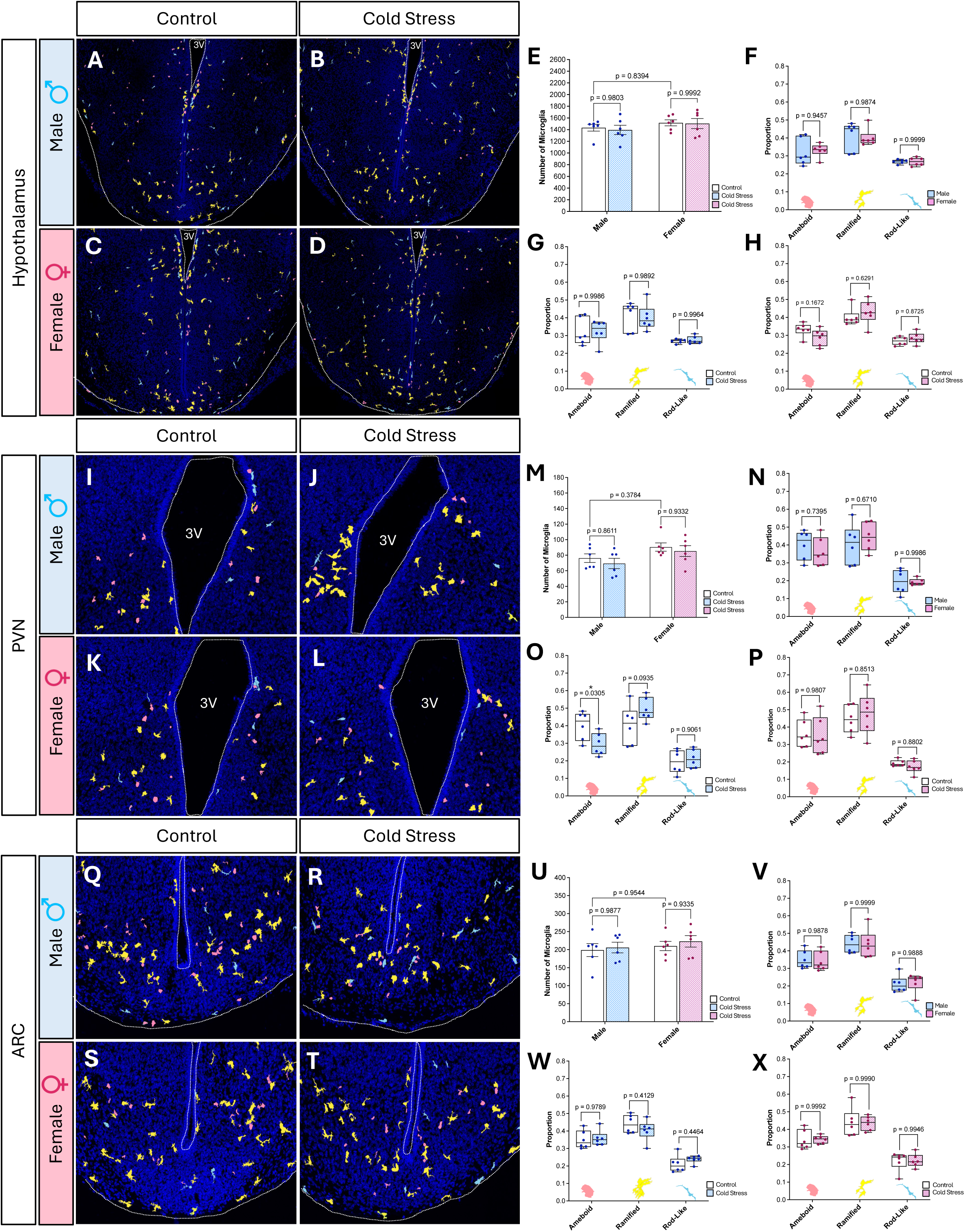
Microglial cluster prevalence in the E15.5 hypothalamus in response to prenatal maternal cold stress. (A–D) Representative images of the E15.5 hypothalamus with ameboid (pink), ramified (yellow), and rod-like (blue) microglia pseudo–colored by cluster. (E) Total number of microglia in the E15.5 hypothalamus (treatment, p=0.7212; sex, p=0.1906; treatment×sex interaction, p=0.8587). (F) Proportion of each microglial cluster across the E15.5 control male and female hypothalamus (cluster, p<0.0001; sex, p=0.9326; cluster×sex interaction, p=0.8496). (G–H) Proportion of each microglial cluster across the E15.5 control and cold stress male (G; cluster, p<0.0001; treatment, p=0.9864; cluster×treatment interaction, p=0.9345) and female (H; cluster, p<0.0001; treatment, p=0.9633; cluster×treatment interaction, p=0.0750) hypothalamus. (I–L) Representative images of the E15.5 PVN. (M) Total number of microglia in the E15.5 PVN (treatment, p=0.3423; sex, p=0.0232; treatment×sex interaction, p=0.8939). (N) Proportion of each microglial cluster in the E15.5 control male and female PVN (cluster, p<0.0001; sex, p=0.8867; cluster×sex interaction, p=0.3937). (O–P) Proportion of each microglial cluster in the E15.5 control and cold stress male (O; cluster, p<0.0001; treatment, p=0.9079; cluster×treatment interaction, p=0.0031) and female (P; cluster, p<0.0001; treatment, p=0.9325; cluster×treatment interaction, p=0.5850) PVN. (Q–T) Representative images of the ARC. (U) Total number of microglia in the E15.5 ARC (treatment, p=0.5238; sex, p=0.3701; treatment×sex interaction, p=0.8530). (V) Proportion of each microglial cluster in the E15.5 control male and female ARC (cluster, p<0.0001; sex, p=0.9617; cluster×sex interaction, p=0.9208). (W–X) Proportion of each microglial cluster in the E15.5 control and cold stress male (W; cluster, p<0.0001; treatment, p=0.9581; cluster×treatment interaction, p=0.1438) and female (X; cluster, p<0.0001; treatment, p=0.9153; cluster×treatment interaction, p=0.9664) ARC. (E, M, U) Data represented as means ± SEM and analyzed using two-way ANOVA with Tukey-adjusted post hoc comparisons. (F–H, N–P, V–X) Data represented as box-and-whisker plots showing the median, interquartile range, and minimum–maximum values and analyzed by a linear mixed-effects model with Sidak-adjusted post hoc comparisons. Each dot represents an individual embryo (n=6), which were collected from 3 independent litters.

Subsequent analysis examined changes in microglial cluster proportions in response to prenatal maternal cold stress in the E15.5 hypothalamus. In males, prenatal maternal cold stress exposure did not alter microglial cluster proportions when the hypothalamus was assessed as a whole (Figure 3G) or when each of the 12 sections spanning the entirety of the hypothalamus were examined separately (Figure S1B, Table S3). Similar to what was observed in males, prenatal maternal cold stress exposure did not significantly alter microglial cluster proportions in females when the hypothalamus was assessed as a whole (Figure 3H). However, when each of the 12 sections spanning the entirety of the hypothalamus were examined separately in females, prenatal maternal cold stress exposure drove a redistribution of microglial clusters in section 3 (Figure S1C, Table S4; cluster×treatment interaction, χ²(2)=15.9706, p=0.0003). This was marked by a significant decrease in the proportion of ameboid microglia (Figure S1C; p=0.0165), and a significant increase in the proportion of ramified microglia (Figure S1C; p=0.0126). No other hypothalamic sections showed differences in microglial cluster proportions between E15.5 control and prenatal maternal cold stress female embryos (Figure S1C, Table S4); however, a significant effect of cluster was observed in both males (Figure 3G; cluster, χ²(2)=35.3002, p<0.0001; Table S3) and females (Figure 3H; cluster, χ²(2)=74.0887, p<0.0001; Table S4).

Given that each hypothalamic section was analyzed independently in our section-by-section analysis, section was not incorporated into the omnibus model or the post-hoc corrections, which may inflate the Type I error rate. Therefore, the changes observed in females in response to prenatal maternal cold stress exposure for section 3 (Figure S1C; Table S4) should be regarded as preliminary. To address this, we re-ran the analysis with section included as a factor. Consistent with our previous analysis, no significant basal differences were detected between control male and female embryos (Figure S2A; Table S5) or between control and prenatal maternal cold stress male embryos (Figure S2B; Table S6). In females, we observed a significant effect of prenatal maternal cold stress on microglial cluster (Figure S2C; Table S7; cluster×treatment interaction, χ²(2)=13.9454, p=0.0009), suggesting that exposure to prenatal maternal cold stress may drive a redistribution of microglial morphology in females. However, no pairwise comparison between control and prenatal maternal cold stress females reached significance (Figure S2C; Table S7). Interestingly, we additionally observed a significant interaction between cluster and hypothalamic section across our control male and female embryos (Figure S2A; Table S5; cluster×section interaction, χ²(22)=51.1091, p=0.0004), as well as in control and prenatal maternal cold stress males (Figure S2B; Table S6; cluster×section interaction, χ²(22)=41.5338, p=0.0071) and females (Figure S2C; Table S7; cluster×section interaction, χ²(22)=51.4863, p=0.0004), highlighting a need for investigation across individual hypothalamic nuclei to better understand how prenatal maternal cold stress may impact microglia in a sex and regionally specific manner.

As we observed regional differences in microglial cluster proportions under control conditions (Figure 2), and prior findings identified sex-specific and microglia-dependent changes in the PVN in response to prenatal maternal cold stress exposure (1), we next assessed microglial morphology in the E15.5 PVN (Figure 3I–L). The number of microglia in the PVN appeared to differ between sexes; however, prenatal maternal cold stress exposure did not affect PVN microglial numbers (Figure 3M; sex, F_(1,20)_=6.047, p=0.0232). While an overall sex difference in the number of microglia in the PVN was detected, where females generally appeared to have higher numbers of microglia, this could not be attributed to any specific group difference following post-hoc analyses (Figure 3M, Table S8). To examine if sex differences in microglial morphology exist in the PVN, we first compared microglial cluster prevalence in the E15.5 male and female PVN under control conditions. Consistent with previous analysis, we observed a significant effect of cluster; however, there was no impact of sex on microglial cluster proportions in the PVN under control conditions (Figure 3N; cluster, χ²(2)=68.9027, p<0.0001). We next examined if microglial cluster proportions changed in response to prenatal maternal cold stress in the E15.5 PVN. In males, prenatal maternal cold stress exposure altered PVN microglial cluster proportions (Figure 3O; cluster×treatment interaction, χ²(2)=11.5224, p=0.0031), with a significant reduction in the ameboid microglial cluster (Figure 3O; p=0.0305), a trend towards an increase in the ramified microglial cluster (Figure 3O; p=0.0935), and no significant change in the rod-like microglial cluster (Figure 3O). In contrast to what was observed in males, prenatal maternal cold stress exposure did not alter PVN microglial cluster proportions in females (Figure 3P). Consistent with prior analysis, a significant effect of cluster was observed in both males (Figure 3O; cluster, χ²(2)=70.9175, p<0.0001) and females (Figure 3P; cluster, χ²(2)=84.8297, p<0.0001).

As the ARC has an established role in satiety and metabolism (84), and prior findings identified altered weight gain in offspring in response to prenatal maternal cold stress exposure (1), we next assessed microglial morphology in the E15.5 ARC (Figure 3Q–T). The number of microglial cells in the ARC did not differ between sexes or in response to prenatal maternal cold stress exposure (Figure 3U, Table S9). To examine if sex differences in microglial morphology exist in the ARC, we first compared microglial cluster prevalence in the E15.5 male and female ARC under control conditions. Consistent with previous analysis, we observed a significant effect of cluster; however, there was no impact of sex on microglial cluster proportions in the ARC under control conditions (Figure 3V; cluster, χ²(2)=104.7325, p<0.0001). We next examined if microglial cluster proportions changed in response to prenatal maternal cold stress in the E15.5 ARC. In males, prenatal maternal cold stress exposure did not alter microglial cluster proportions in the ARC (Figure 3W). Similar to what was observed in males, exposure to prenatal maternal cold stress did not alter microglial cluster proportions in the ARC of females (Figure 3X). Again, a significant effect of microglial cluster was observed in both males (Figure 3W; cluster χ²(2)=124.8469, p<0.0001) and females (Figure 3X; cluster χ²(2)=120.0564, p<0.0001).

Collectively, these findings demonstrate that microglial morphology displays regional differences within the developing fetal hypothalamus, with prenatal maternal cold stress exposure driving a redistribution of microglial morphologies within the PVN of male embryos. These results underscore the importance of considering sex and region in analysis of microglial morphology in the developing brain.

### Supervised clustering reveals a continuum of morphological clusters and unique responses to prenatal maternal cold stress in the fetal male and female PVN

Given there are a wide array of clustering approaches and the differences they can show in assessment of microglial morphology (82,83), an additional clustering method was employed to evaluate microglial morphology in the E15.5 PVN—a region that displayed unique responses to prenatal maternal cold stress across males and females (Figure 3I-P). This approach employed MorphoGlia, a software package for microglial morphology developed by Maya-Arteaga et al. (2024), which utilizes supervised feature selection, Uniform Manifold Approximation and

Projection (UMAP) dimensionality reduction, and Hierarchical Density-Based Spatial Clustering of Applications with Noise (HDBSCAN) clustering (85). Microglial cells were extracted and skeletonized within the MorphoGlia application, yielding 32 distinct morphological features, many of which were highly correlated (Figure 4A). Examination of individual features in the E15.5 male PVN revealed that control male embryos were characterized by greater soma circularity, cell solidity, and fractal dimension—hallmarks of a more ameboid morphology, whereas prenatal maternal cold stress male embryos showed greater branch number, Sholl crossings, and cell area, indicative of a more ramified morphology (Figure 4B). These differences were not observed in the E15.5 female PVN, where control and prenatal maternal cold stress female embryos exhibited relatively similar morphological attributes (Figure 4C).

**Figure 4.**
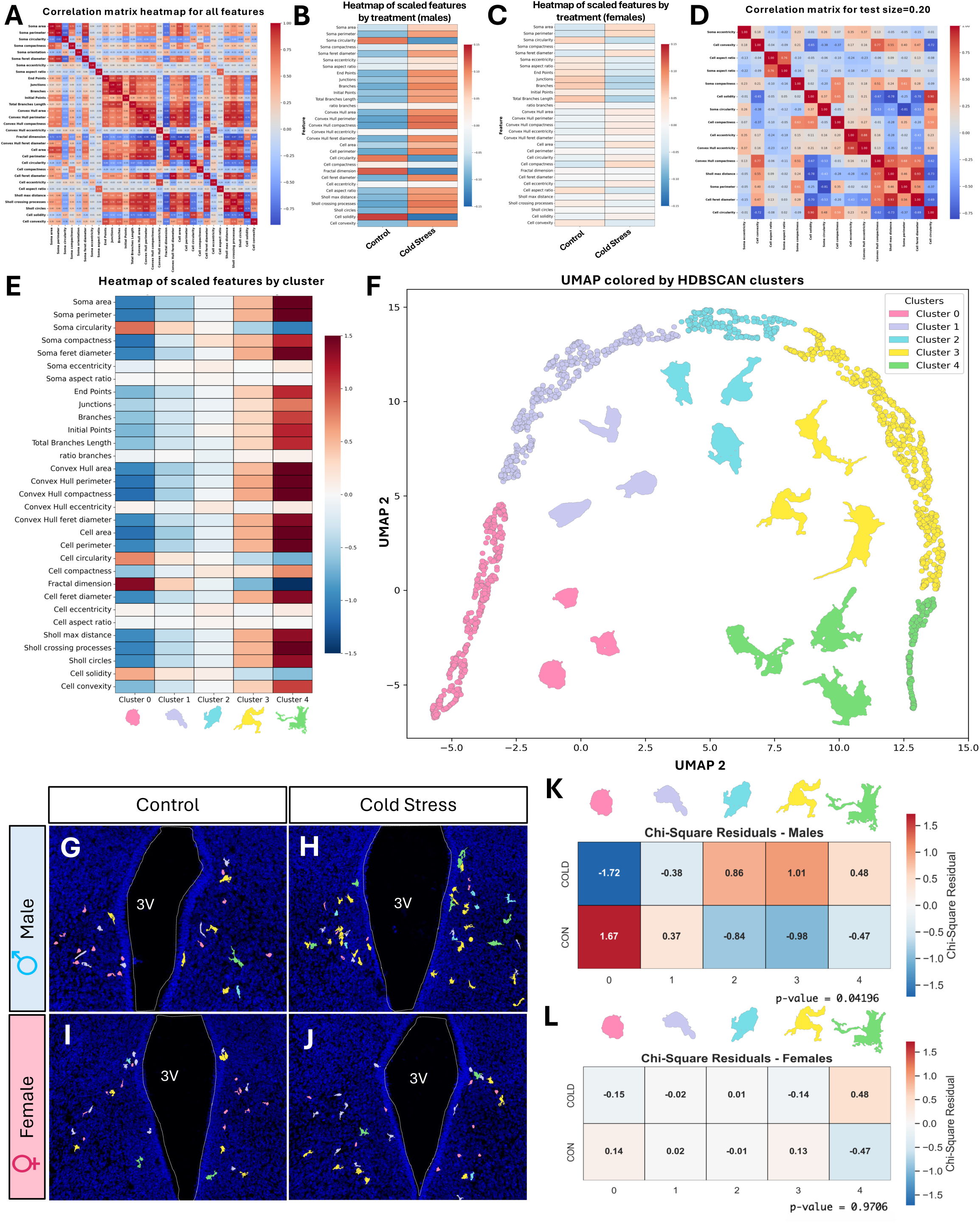
Morphological clustering and quantification using supervised feature selection in the E15.5 PVN. (A) Heatmap of pairwise Pearson correlations among all morphological features extracted. (B–C) Heatmaps comparing morphological feature distributions between control and cold stress male (B) and female (C) microglia in the E15.5 PVN. (D) Heatmap of pairwise Pearson correlations among morphological features selected using random feature elimination. (E) Heatmap summarizing the relationship between the 32 individual morphological features extracted from each microglia and cluster identity. (F) UMAP embedding of single-cell morphological features with cluster assignments, showing five distinct microglial clusters. (G–J) Representative images with microglia pseudo–colored by cluster identity in the E15.5 control male (G), cold stress male (H), control female (I), and cold stress female (J) PVN. (K) Pearson chi-squared test comparing microglial cluster proportions in the E15.5 control and cold stress male PVN (χ²(4)=9.91, p=0.04196). (L) Pearson chi-squared test comparing microglial cluster proportions in the E15.5 control and cold stress female PVN (χ²(4)=0.53, p=0.9706). Microglia were extracted from n=6 embryos per sex/treatment from 3 independent litters.

Given these observations and the effects of prenatal maternal cold stress on microglial morphology identified in our previous clustering analysis (Figure 3I-P), treatment was set as the contrast of interest (i.e., control vs. cold stress), and a random forest classifier was applied to identify the 16 most informative morphological features distinguishing control and prenatal maternal cold stress groups (Figure 4D). UMAP dimensionality reduction followed by HDBSCAN clustering of these features revealed five distinct clusters spanning a morphological gradient from ameboid (Cluster 0) to a more ramified (Cluster 4) morphology (Figure 4E, F).

Next, we were interested in visualizing whether group differences in microglial cluster prevalence existed in the E15.5 control and prenatal maternal cold stress male and female PVN (Figure 4G–J). Similar to what we observed using the MicrogliaMorphology pipeline established by Kim et al. (2024), prenatal maternal cold stress drove a shift away from ameboid and toward a more ramified microglial morphology in the E15.5 PVN of males (Figure 4K; χ²(4)=9.91, p=0.04196), driven primarily by a reduction in the ameboid cluster (Cluster 0) and a redistributed increase across the more ramified clusters (Cluster 2 and 3). In contrast, microglial cluster distribution in the E15.5 PVN appeared consistent between control and prenatal maternal cold stress female embryos, with no significant changes observed in any microglial clusters (Figure 4L).

Together, these data strengthen the findings generated using the MicrogliaMorphology pipeline established by Kim et al. (2024), where prenatal maternal cold stress exposure was shown to drive a redistribution of microglial morphology in the PVN of male embryos.

Furthermore, by utilizing two independent pipelines for the assessment of microglia morphology, we illustrate that whether microglial morphology is captured along a continuous spectrum or in a more discrete categorical manner, both approaches ultimately yield similar findings and highlight the robustness of the underlying changes in microglial morphology.

### Exposure to prenatal maternal cold stress drives unique changes in microglia–AVP neuron interactions in the fetal male and female PVN

Considering that exposure to prenatal maternal cold stress has previously been shown to reduce OXT neuron numbers in the E15.5 PVN of male embryos and drive microglia-dependent social deficits in male offspring (1), we next wanted to determine if the shift in microglial morphology we observed in the fetal PVN had functional implications. We began our investigation by examining microglia-AVP neuron contacts in the E15.5 PVN, as AVP neurons have been implicated in social deficits (61,63), yet their numbers were unchanged in response prenatal maternal cold stress exposure (1). Accordingly, changes in microglia–AVP neuron contact may represent an upstream target through which the developmental disruptions in social behaviors may occur following prenatal maternal cold stress exposure. Microglial contact with AVP neurons was examined across two categories: touching events and wrapping events. Touching events were defined visually by microglia–AVP neuron soma contacts without significant soma overlap (Figure 5A) or with one to two microglial branches overlapping in the absence of soma-to-soma contact (Figure 5B). Wrapping events were defined visually as cases involving soma overlap that also included branch interactions (Figure 5C) or significant soma interactions (Figure 5D). Microglia–AVP neuron interactions were then mapped back to their original morphological clusters to assess differences in the number and type of interactions between experimental groups and across clusters. Imaris 3D renderings were used to visualize touching (Figure 5E) and wrapping (Figure 5F) events.

**Figure 5.**
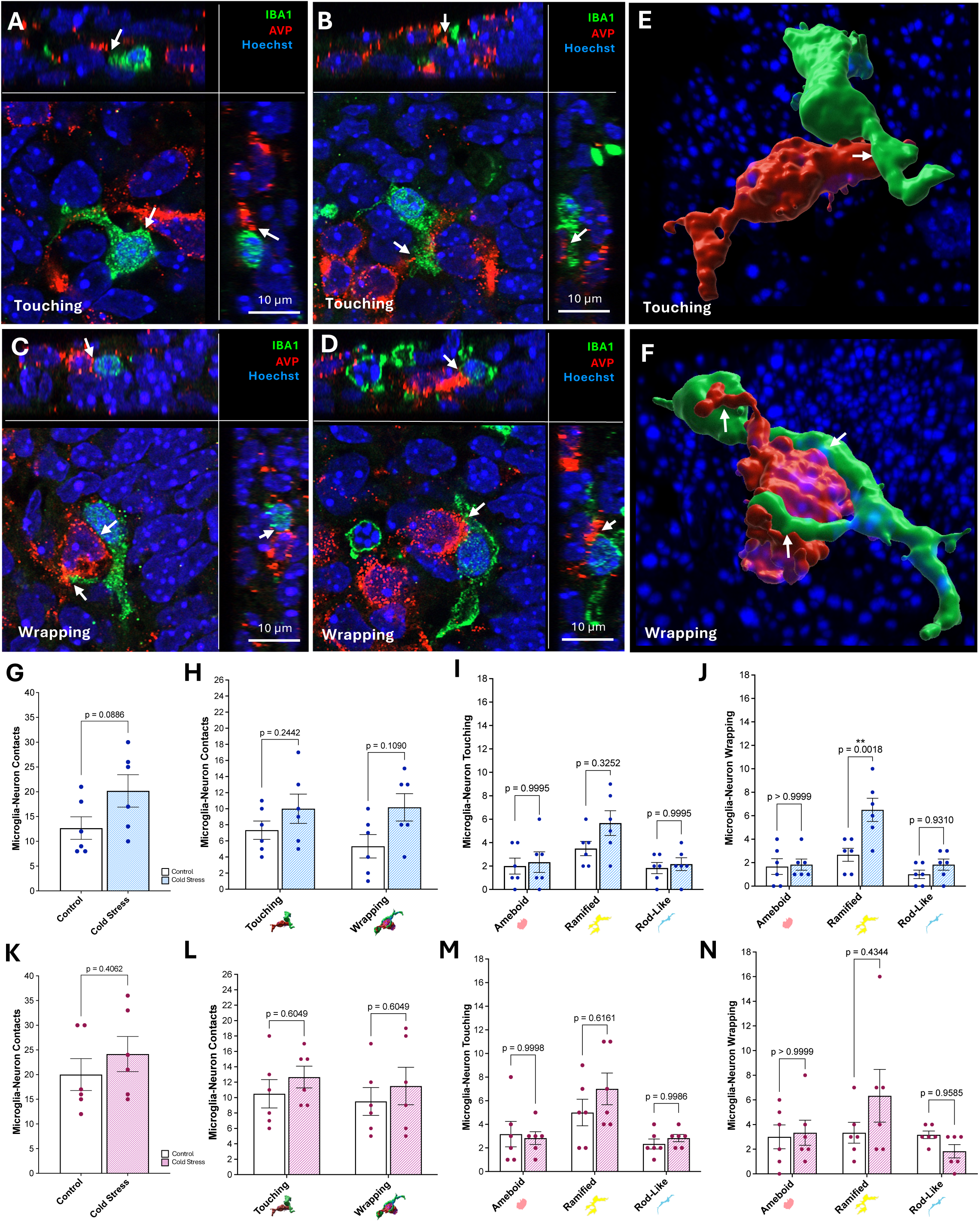
Microglia–AVP neuron contacts in the E15.5 PVN. (A–D) Representative images of microglia–AVP neuron touching (A, B) and wrapping (C, D) events in the E15.5 PVN. (E–F) Representative *Imaris* reconstructions of microglia (green) and AVP neurons (red), showing touching (E) and wrapping (F) events in the E15.5 PVN. (A–F) White arrows mark sites of contact. (G–H) Total microglia–AVP neuron contacts (G) and contacts by event type (touching vs. wrapping) (H) in the E15.5 control and cold stress male PVN. (I) Microglia–AVP neuron touching events by cluster (ameboid vs. ramified vs. rod-like) in the E15.5 control and cold stress male PVN (cluster, p=0.0020; treatment, p=0.1271; cluster×treatment interaction, p=0.3691). (J) Microglia–AVP neuron wrapping events by cluster in the E15.5 control and cold stress male PVN (cluster, p<0.0001; treatment, p=0.0035; cluster×treatment interaction, p=0.0142). The number of ramified microglia–AVP neuron wrapping events is significantly higher in cold stress males (p=0.0018). (K–L) Total microglia–AVP neuron contacts (K) and contacts by event type (L) in the E15.5 control and cold stress female PVN. (M) Microglia–AVP neuron touching events by cluster in the E15.5 control and cold stress female PVN (cluster, p=0.0010; treatment, p=0.3303; cluster×treatment interaction, p=0.4273). (N) Microglia–AVP neuron wrapping events by cluster in the E15.5 control and cold stress female PVN (cluster, p=0.1214; treatment, p=0.4754; cluster×treatment interaction, p=0.1713). (G–N) Data are represented as means ± SEM and were analyzed by a Student’s t-test (G, K), Student’s t-test with Holm-Sidak adjustment for multiple comparisons (H, L) or two-way ANOVA with Tukey-adjusted post hoc comparisons (I, J, M, N). Each dot represents an individual embryo (n=6), which were collected from 3 independent litters.

To begin, the number of microglia–AVP neuron interactions were compared across the E15.5 control male and female PVN, starting across all microglia–AVP neuron contacts (touching and wrapping events combined). A trend toward increased microglia–AVP neuron contacts was observed in females as compared to males, although this did not reach significance (Figure S3A; p=0.0934). Stratification by contact type (i.e., touching versus wrapping) showed no significant differences in microglia–AVP neuron touching or wrapping events between control male and female embryos (Figure S3B). To better assess whether the previously observed shifts in microglial morphology had functional implications, we further stratified touching and wrapping events by microglial cluster (i.e., ameboid, ramified, and rod-like). We observed no differences in microglia–AVP neuron touching events in any cluster between control male and female embryos (Figure S3C, Table S10). Interestingly, when stratified by cluster, we did observe a significant effect of sex on the number of microglia–AVP neuron wrapping events in control embryos (Figure S3D; Table S11; sex, F_(1,30)_=6.607, p=0.0154), where female embryos appeared to have a greater number of these events at baseline. This suggests that there could be subtle differences in the number of microglia–AVP neuron wrapping events across control male and female embryos; however, this could not be attributed to any specific within-cluster comparison (Figure S3D, Table S11). A significant effect of cluster was also observed across control embryos for touching events (Figure S3C; Table S10; cluster, F_(2,30)_=4.193, p=0.0248) but interestingly was not observed for wrapping events (Figure S3D, Table S11).

Next the effect of prenatal maternal cold stress was assessed in male and female embryos, starting with all microglia–AVP neuron contacts within the E15.5 male PVN. A trend toward increased microglia–AVP neuron contacts was observed in males in response to prenatal maternal cold stress exposure, although this did not reach significance (Figure 5G; p=0.0886).

Similarly, stratification by contact type revealed no significant differences in microglia–AVP neuron touching or wrapping events in the E15.5 control and prenatal maternal cold stress PVN of male embryos (Figure 5H). To better assess whether the previously observed shifts in microglial morphology had functional implications, we again stratified touching and wrapping events by microglial cluster. Although prenatal maternal cold stress did not alter microglia–AVP neuron touching events in any cluster (Figure 5I, Table S12), microglia–AVP neuron wrapping events were significantly altered by prenatal maternal cold stress exposure in a cluster-dependent manner (Figure 5J, Table S13; treatment, F_(1,30)_=10.04, p=0.0035; cluster×treatment interaction, F_(2,30)_=4.916, p=0.0142). Specifically, the number of ramified microglia wrapping around AVP neurons increased in males in response to prenatal maternal cold stress exposure (Figure 5J; p=0.0018), with no significant change to ameboid or rod-like microglia–AVP neuron wrapping events (Figure 5J). Consistent with prior analyses, a significant effect of cluster was observed for both touching (Figure 5I, Table S12; cluster, F_(2,30)_=7.692, p=0.0020) and wrapping (Figure 5J, Table S13; cluster, F_(2,30)_=15.61, p<0.0001) events.

In females, the number of microglia–AVP neuron contacts in the PVN did not differ significantly between E15.5 control and prenatal maternal cold stress embryos (Figure 5K). Moreover, stratification by contact type revealed no significant differences in microglia–AVP neuron touching or wrapping events between the E15.5 control and prenatal maternal cold stress PVN of female embryos (Figure 5L). Furthermore, cluster-stratified analysis demonstrated that prenatal maternal cold stress exposure did not alter microglia–AVP neuron touching events (Figure 5M, Table S14) or wrapping events (Figure 5N, Table S15) in the E15.5 PVN of females for any cluster. Consistent with prior analyses, a significant effect of cluster was observed for microglia–AVP neuron touching events in the E15.5 PVN of female embryos (Figure 5M, Table S14; cluster, F_(2,30)_=8.696, p=0.0010). Notably, no significant effect of cluster was observed for microglia–AVP neuron wrapping events in the E15.5 PVN of female embryos (Figure 5N, Table S15), suggesting a possible difference in the distribution of microglial clusters involved in AVP neuron wrapping events in females as compared with other analyses.

Given the changes in ramified microglia–AVP neuron wrapping events seen in male embryos in response to prenatal maternal cold stress exposure, we next assessed whether the proportion of ramified microglia engaged in AVP neuron contacts was similarly altered. While only ramified microglia–AVP neuron wrapping events reached significance, all contact types (i.e., touching and wrapping events) were included in this analysis to capture the full extent of ramified microglia–AVP neuron engagement. Interestingly, exposure to prenatal maternal cold stress drove a significant increase in the proportion of ramified microglia in contact with AVP neurons in the E15.5 PVN of male embryos (Figure S4A; p=0.0133). This was unique to male embryos, as prenatal maternal cold stress exposure did not alter the proportion of ramified microglia in contact with AVP neurons in the E15.5 PVN of female embryos (Figure S4B).

Taken together, the observed male-specific increase in ramified microglia–AVP neuron interactions (i.e., wrapping events) in response to prenatal maternal cold stress exposure is consistent with the morphological transition away from an ameboid and toward a ramified microglial morphology seen in the fetal PVN of males, suggesting that this morphological shift may correspond to broader changes in microglial behaviors.

### Prenatal maternal cold stress exposure drives unique changes in hypothalamic microglial engulfment behavior in male and female embryos

Considering that microglial phagocytosis plays a prominent role in neurodevelopment (39,42,86), and there is evidence that microglial phagocytic activity is altered in the fetal hypothalamus in response to prenatal maternal perturbations (49), we next examined the impact of prenatal maternal cold stress on microglial phagocytic activity in the developing hypothalamus. To begin, microglial phagocytic activity was assessed in the E15.5 PVN, where microglia were classified as phagocytic based on the presence of at least one phagocytic cup containing Hoechst+ nuclear material with clear circular morphology (Figure 6A–D). At baseline, no differences were observed in microglial phagocytic activity between control male and female embryos (Figure 6E), and cluster-stratified analysis (i.e., ameboid vs. ramified vs. rod-like) further demonstrated no differences across any microglial clusters between control male and female embryos (Figure 6F; Table S16). Next, we assessed changes in microglial phagocytosis in response to prenatal maternal cold stress. Although the number of phagocytic microglia in the PVN did not differ between E15.5 control and prenatal maternal cold stress male embryos (Figure 6G), cluster-stratified analysis demonstrated that prenatal maternal cold stress exposure resulted in a cluster-dependent change in the number of phagocytic microglia in the E15.5 PVN of males (Figure 6H, Table S17; cluster×treatment interaction, F_(2,30)_=4.259, p=0.0235). In males, prenatal maternal cold stress exposure increased the number of phagocytic ramified microglia (Figure 6H; p=0.0254), with no significant changes observed in phagocytic ameboid or rod-like microglia (Figure 6H). In females, the number of phagocytic microglia in the PVN did not differ between E15.5 control and prenatal maternal cold stress embryos (Figure 6I), even when cluster-stratified analysis was performed (Figure 6J, Table S18). Consistent with prior analyses, a significant effect of cluster was observed in both males (Figure 6H, Table S17; cluster, F_(2,30)_=27.44, p<0.0001) and females (Figure 6J, Table S18; cluster, F_(2,30)_=21.67, p<0.0001).

**Figure 6.**
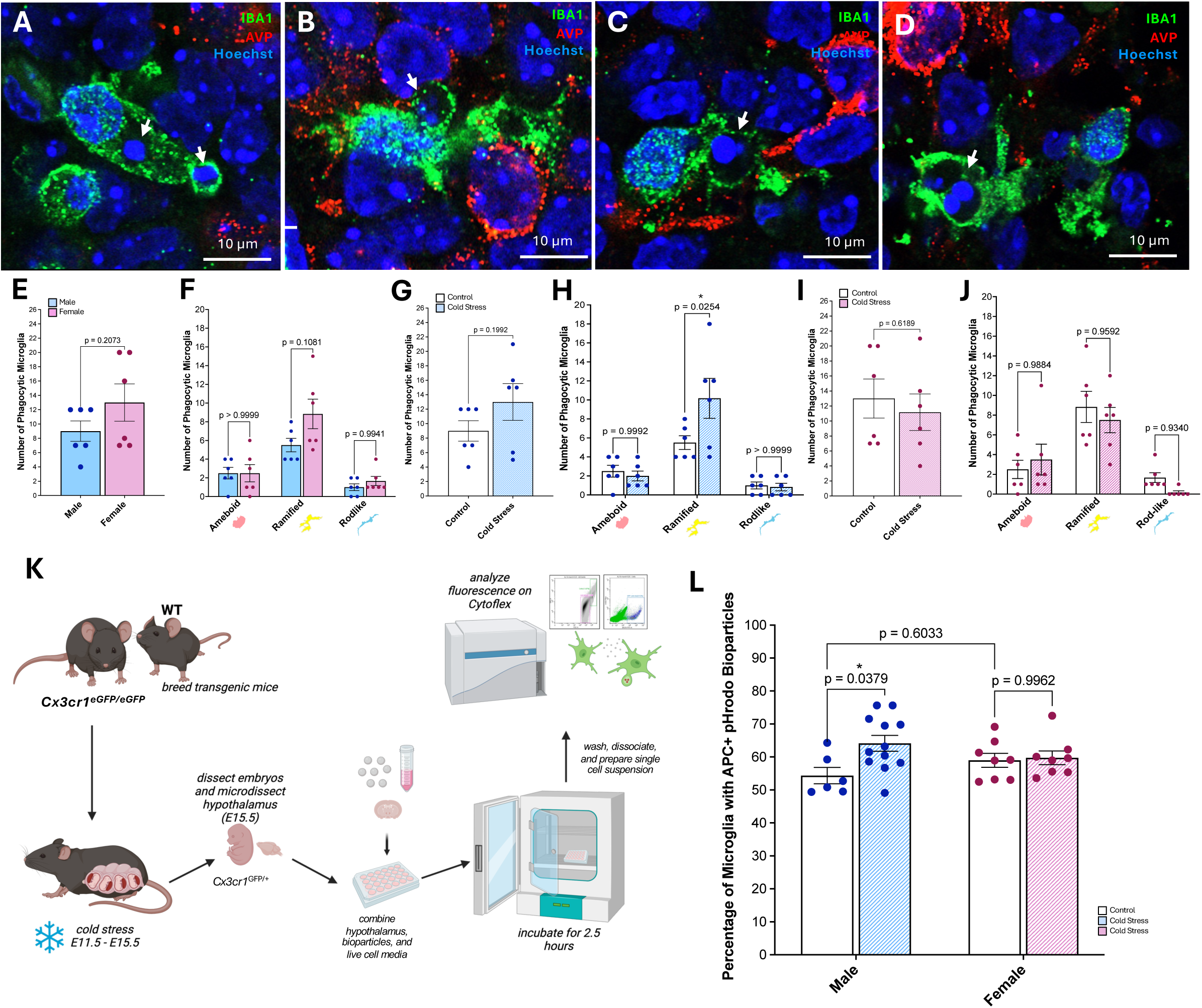
Microglial phagocytosis in the E15.5 hypothalamus. (A–D) Representative images of phagocytic microglia in the E15.5 PVN. White arrows mark phagocytic events. (E) Number of phagocytic microglia in the E15.5 control male and female PVN (p=0.2073). (F) Number of phagocytic microglia by cluster (ameboid vs. ramified vs. rod-like) in the E15.5 control male and female PVN (cluster, p<0.0001; sex, p=0.0726; cluster×sex interaction, p=0.1504). (G) Number of phagocytic microglia in the E15.5 control and cold stress male PVN (p=0.1992). (H) Number of phagocytic microglia by cluster in the E15.5 control and cold stress male PVN (cluster, p<0.0001; treatment, p=0.1097; cluster×treatment interaction, p=0.0235). The number of ramified phagocytic microglia is significantly higher in cold stress males (p=0.0254). (I) Number of phagocytic microglia in the E15.5 control and cold stress female PVN (p=0.6189). (J) Number of phagocytic microglia by cluster in the E15.5 control and cold stress female PVN (cluster, p<0.0001; treatment, p=0.5143; cluster×treatment interaction, p=0.4767). (K) Schematic outlining flow cytometry bioparticle assay for assessing microglial phagocytic capacity in the E15.5 hypothalamus. (L) Percentage of microglia with engulfed bioparticles in the E15.5 control and cold stress male and female hypothalamus (treatment, p=0.0385; sex, p=0.9610; treatment×sex interaction, p=0.0736). The percentage of microglia with engulfed bioparticles is significantly higher in cold stress males compared to control males (p=0.0379). (E–J, L) Data are represented as means ± SEM and were analyzed using a Student’s t-test (E, G, I) or two-way ANOVA with Tukey-adjusted post hoc comparisons (F, H, J, L). Each dot represents an individual embryo (n=6-12), which were collected from 3-4 independent litters.

Next, we assessed if the proportion of ramified microglia exhibiting phagocytic activity was altered by prenatal maternal cold stress in the E15.5 PVN. No significant change in the proportion of ramified phagocytic microglia was observed in the E15.5 PVN between control and prenatal maternal cold stress males or females (Figure S4C-D). This suggests that while prenatal maternal cold stress increases ramified phagocytic microglia in the E15.5 PVN of male embryos, this increase is proportional to the overall expansion of the ramified population seen in Figure 3, rather than reflecting a preferential recruitment of this cluster towards increased phagocytic activity.

Given the increase in microglial phagocytic activity observed in the PVN of male embryos in response to prenatal maternal cold stress exposure, we sought to extend this analysis to the broader hypothalamus using a complementary approach. To achieve this, hypothalamic tissue was micro-dissected from E15.5 control and prenatal maternal cold stress male and female *Cx3cr1^GFP/+^* transgenic embryos and microglial phagocytosis was assessed using pHrodo—a pH-sensitive bioparticle conjugate that is non-fluorescent at neutral extracellular pH but fluoresces red upon internalization into the acidic phagolysosomal environment (87), allowing for quantification of internalized particles via flow cytometry. pHrodo bioparticles have been validated for quantifying phagocytic activity across several cell types, including microglia, macrophages, and granulocytes (87–89). Here, we quantified the percentage of microglia (GFP+) exhibiting red fluorescence (APC+) as our measure of phagocytic activity (Figure 6K, S5A–C), observing a significant impact of prenatal maternal cold stress on microglial phagocytic activity in the E15.5 hypothalamus (Figure 6L, Table S19; treatment, F_(1,30)_=4.683, p=0.0385; treatment×sex interaction, F_(1,30)_=3.437, p=0.0736). Consistent with a trend toward a sex-dependent effect, only microglia collected from prenatal maternal cold stress male hypothalami showed a significant increase in pHrodo bioparticle uptake relative to control males (Figure 6L; p=0.0379), while bioparticle uptake did not differ between microglia analyzed from control and prenatal maternal cold stress female hypothalami (Figure 6L).

Together, these findings indicate that exposure to prenatal maternal cold stress drives unique cluster-specific changes in microglial phagocytic activity in the PVN of male and female embryos, and perhaps more broadly across the developing hypothalamus as indicated by the engulfment of pHrodo bioparticles.

## Discussion

Maternal stress during pregnancy is widely associated with long-term changes in offspring neuroanatomy, physiology, and behavior. These effects often show sex differences (23,24,26–31), suggesting that exposure to prenatal maternal stress may influence vulnerability to NDDs in a sex-dependent manner. Fetal microglia are positioned to contribute to these outcomes as they are highly responsive to the intrauterine environment and actively shape neural circuits across development (1,34,38–43,49). Despite this, our understanding of how prenatal maternal stress alters fetal microglia at the cellular level remains largely unexplored, particularly within the hypothalamus. Additionally, while microglial morphology is commonly interpreted as a proxy for cellular state (82), few studies have directly linked fetal microglial morphology with functional behaviors such as interactions with neurons and phagocytosis. To address this gap, we adapted two pipelines, namely MicrogliaMorphology (79) and MorphoGlia (85) to assess fetal microglial morphology in order to examine how prenatal maternal cold stress impacts microglial morphology and functionally determine whether changes in morphology could be linked to altered cellular interactions and/or phagocytic behaviours in the E15.5 hypothalamus of male and female embryos. Although we did not observe sex differences in microglial morphology between control males and females, in response to prenatal maternal cold stress, microglia displayed sex- and region-specific changes. In the E15.5 male PVN, exposure to prenatal maternal cold stress shifted microglia from an ameboid morphology towards a more ramified morphology. This was coupled with increased interactions between ramified microglia and AVP neurons, as well as an increase in the number of ramified microglia displaying phagocytic activity. These findings were complemented by analysis of the whole hypothalamus using flow cytometry, which similarly showed increased microglial engulfment of pHrodo bioparticles in E15.5 male embryos only.

Together, these findings indicate that exposure to prenatal maternal stress causes unique changes in microglial morphology and function in the fetal hypothalamus of male and female embryos, and builds on evidence that male microglia may be uniquely impacted by this prenatal maternal stress model (1).

Research suggests that microglial morphology can differ by brain region, sex, and developmental time-point (53,54). In line with this, we observed heterogeneity across the fetal hypothalamus and within specific nuclei, with a higher proportion of rod-like microglia seen in the whole hypothalamus as compared to the PVN and ARC. While the functional significance of rod-like microglia is incompletely understood, this morphology has been associated with microglial polarization, migration, alignment along neuronal processes such as dendrites and axons, and disease (90–92). This uneven distribution of microglia across different hypothalamic nuclei may reflect differing microglial functions and is not unique to the hypothalamus. For example, in the basal ganglia, microglia have been shown to differ between neighboring nuclei in density, branching morphology, and lysosome content beginning in the second postnatal week (93). Given that the hypothalamus is among the earliest brain regions to be colonized by microglia and to begin functional differentiation (34,65,94), regional heterogeneity may also emerge earlier, which is consistent with our observations at E15.5.

Except for the PVN, where female embryos appeared to have higher numbers of microglia in comparison to male embryos, we did not observe baseline sex differences in microglial counts or morphology across the PVN, ARC or whole hypothalamus in any post-hoc comparisons in our static analysis. This is consistent with prior work where limited sex differences have been reported for microglia in the fetal brain under homeostatic conditions (53,95). However, differences may emerge later, as research shows that sex-divergent microglial phenotypes increase during the postnatal period (53), and relevant to this study, research also indicates that fetal microglia show both region- and sex-specific responses to prenatal maternal stressors (1,9,12,49,53,55,71,95). While we observed no effect of prenatal maternal cold stress on microglial morphology in either sex when the hypothalamus was examined as a whole, sub-regional analysis across our 12 rostro-caudal hypothalamic sections revealed a shift in microglial morphology in section 3 of female embryos, marked by a reduction in the proportion of ameboid microglia and an increase in the proportion of ramified microglia. Further exploration of this finding is required as it may indicate a localized response in the anterior hypothalamus or a specific nucleus within this region. Interestingly, when we restricted our analysis to the PVN, a parallel shift emerged in males in response to prenatal maternal cold stress: male embryos showed a significant reduction in the proportion of ameboid microglia and a trend toward an increase in the proportion of ramified microglia. This shift from an ameboid to ramified morphology is intriguing, as ramified microglia are often interpreted as being in a resting or homeostatic state in the adult brain (95,96). However, this interpretation does not translate directly to fetal microglia, which are developmentally distinct and predominantly ameboid even at baseline (53). For instance, in the fetal hypothalamus, prenatal exposure to bisphenol A (BPA) has been shown to increase microglial ramification (49). Additionally, even in adults, increased ramification does not always reflect a homeostatic state. For example, chronic stress has been shown to drive a hyper-ramified phenotype in the rat prefrontal cortex, marked by increased branching complexity and process length (97). Comparable phenotypes have also been reported in the hippocampus and medial prefrontal cortex in mouse models of post-traumatic stress disorder (PTSD) (98). Interestingly, in immune perturbation models, microglia often adopt a reactive hypertrophic morphology, characterized by an enlarged soma and shorter, thicker processes (46,99). Because our clustering approach did not resolve a hyper-ramified or hypertrophic morphology, functionally distinct microglial phenotypes may have been grouped within the ramified cluster. This is particularly relevant in the embryo, where microglia are predominantly ameboid under homeostatic conditions, making reactive and resting states especially difficult to separate by morphology alone.

For these reasons, morphology cannot fully distinguish a resting from a reactive microglial state. Interestingly, in male embryos, exposure to prenatal maternal cold stress also increased both the number and proportion of ramified microglia engaged in wrapping events with AVP neurons, suggesting that this shift in microglial morphology in response to prenatal maternal stress has functional implications. Because both the number and proportion of ramified microglia engaged in wrapping events with AVP neurons increased, a greater fraction of ramified microglia interacted with AVP neurons than would be expected from the rise in ramified microglia alone—indicating preferential engagement of this microglial cluster. This interpretation is informative alongside previous findings that exposure to prenatal maternal cold stress produced a microglia-dependent reduction in OXT neurons, but not AVP neurons, in the E15.5 PVN of male embryos (1). Our findings suggest that although AVP neuron number remains unchanged at E15.5, exposure to prenatal maternal cold stress may still alter local microglial interactions with AVP neurons and could impact AVP neuron numbers at later stages of development and/or AVP neuron physiology. Given that both OXT and AVP neurons are central regulators of social behaviors and have been implicated in ASD-related phenotypes (61), this suggests that prenatal maternal cold stress may affect multiple PVN neuronal populations in distinct ways. This aligns with emerging evidence that microglia in the PVN may interact with neurons in a neuropeptide-specific manner. For example, in the early postnatal PVN, microglia contacting CRH neurons have been shown to engulf fewer excitatory synapses than neighboring microglia not in contact with CRH neurons (100,101), illustrating not only that microglia and neurons communicate, but that the neuropeptide identity of nearby neurons can shape this communication and ultimately microglial behavior.

In addition to altered ramified microglia–AVP neuron wrapping events, exposure to prenatal maternal cold stress also increased the number of ramified phagocytic microglia in the E15.5 PVN of male embryos, which was not observed in females. Notably, while ramified phagocytic microglia counts increased, proportions did not, reflecting the expansion of the ramified microglial cluster in the E15.5 PVN of prenatal maternal cold stress male embryos, rather than preferential recruitment of this microglial cluster towards a phagocytic state. This contrasts with what was observed for microglia–AVP neuron interactions, where both number and proportion increased, and indicates that prenatal maternal cold stress may engage ramified microglia in the fetal PVN differently across these two functional behaviors. However, in the case of phagocytic microglia, an increase in their numbers in response to prenatal maternal cold stress, even in the absence of a change in proportion, would likely still ultimately result in increased phagocytosis in the PVN, which could have detrimental impacts on the developing brain.

Complementing our static analysis of microglial phagocytic activity in the fetal PVN, we also observed an increase in microglial phagocytosis in the whole hypothalamus of male embryos in response to prenatal maternal cold stress when using a pHrodo bioparticle assay. Because the pHrodo bioparticles are exogenous targets (inactivated *E. coli*), the assay is widely used as a quantitative measure of microglial and macrophage phagocytic capacity (87–89). Interpreted in this framework, our results suggest that exposure to prenatal maternal cold stress increases the phagocytic capacity of hypothalamic microglia, but only in male embryos. Given this effect was detected in tissue collected from whole hypothalami, this suggests that the increase in microglial phagocytosis seen in male embryos in response to prenatal maternal cold stress exposure may not be restricted to the PVN. It is important to note that because we included the entire litter for these experiments, and flow cytometry is susceptible to batch effects arising from instrument-related variability such as day-to-day differences in cytometer performance, laser power, and detector sensitivity across separate acquisitions, instrument-related variability cannot be excluded. However, the sex differences we observed across both of our microglial phagocytosis assays parallel other reports showing sex-specific microglial phagocytosis in response to prenatal perturbation. For instance, maternal exposure to high-fat diet drives excess microglial phagocytosis of serotonin neurons in the dorsal raphe nucleus of male embryos, and results in sex-specific behavioral outcomes (102). This is particularly relevant as phagocytosis is essential for shaping neural circuits, and both excessive and deficient synaptic pruning during development have been linked to NDDs (103). For example, insufficient pruning has been associated with ASD-like phenotypes, whereas excessive pruning has been implicated in schizophrenia (104,105). However, this directionality is not absolute and the opposite pattern has also been documented with some ASD-associated models showing excessive microglial pruning of neuronal synapses and neuronal populations (22,102). Accordingly, the changes in phagocytic activity we observed in male embryos across our static and flow cytometry analyses, in both the PVN and whole hypothalamus, may represent one candidate cellular pathway through which prenatal maternal stress could contribute to sex-specific disruptions in neurodevelopmental trajectories.

## Conclusions

This study is the first to adapt MicrogliaMorphology (79) and MorphoGlia (85) pipelines to study fetal microglial morphology and link unique morphological shifts to functional changes in microglial behavior, together helping to further advance our understanding of how prenatal maternal stress reshapes microglial activity within the fetal brain. We provide novel region-specific characterizations of microglial morphology within the fetal hypothalamus of both male and female embryos, establishing unique sex differences in fetal hypothalamic microglia sensitivity to prenatal maternal stress. More broadly, this research identifies fetal hypothalamic microglia as a potential cellular link between perturbation of the intrauterine environment and sex differences in neurodevelopmental outcome. By curating currently available open-source tools for assessing microglial morphology (79,85) to work in the fetal brain, and highlighting the importance of refining your analyses in a region-specific manner in both sexes, these findings provide a foundation for future studies to examine how early-life perturbation of the intrauterine environment impact microglial responsiveness and contribute to later sex-specific changes in circuit development and behavior.

## Methods

All code associated with morphological clustering, data frames with microglial morphology measures, statistical analysis in R, video tutorials, and microglial inclusion/exclusion criteria can be found on GitHub: https://github.com/RosinLabUBC/Microglia-Cold-Stress-Project-AL.git

### Experimental model

All experimental work was conducted in accordance with Biosafety Protocol B21-0133, which received approval from the University of British Columbia. Animal work was carried out in accordance with the guidelines and regulations of the Canadian Council on Animal Care and received prior approval from the University of British Columbia’s Animal Care Committee (protocols A21-0170 and A21-0171).

Male and female CD1 mice (strain code 022, Charles River) were used for all static analyses of microglia, including microglial clustering for morphological assessment and functional analysis (e.g., phagocytic cups and interactions with neurons). To generate heterozygous *Cx3cr1^eGFP^* embryos for flow cytometry experiments, homozygous *Cx3cr1^eGFP/eGFP^* mice (*B6.129P2(Cg)-Cx3cr1^tm1Litt^/J*; The Jackson Laboratory 005582), which express green fluorescent protein (eGFP) under control of the endogenous *Cx3cr1* locus, were crossed with C57BL/6J mice. Breeding pairs consisted of either homozygous *Cx3cr1^eGFP/eGFP^*males with wild-type (WT) females or WT males with homozygous *Cx3cr1^eGFP/eGFP^* females, resulting in exclusively heterozygous *Cx3cr1^eGFP/+^* embryos. All experiments were blinded to sex and/or treatment.

### Mouse handling and maternal cold stress paradigm

Vaginal plugs were checked daily in the morning following the setup of a breeding pair. The day of plug detection was designated embryonic day 0.5 (E0.5). Pregnant mice were housed individually and received standard chow (LabDiet PicoLab Rodent Diet 20) and water *ad libitum* during pregnancy. Pregnant mice were randomly assigned to control or cold stress conditions.

Prenatal maternal cold stress was conducted by placing pregnant mice in a 4°C walk-in refrigerator for 30 min daily from E11.5 to E15.5, during which time pregnant mice remained in their home cages with *ad libitum* access to standard chow (LabDiet PicoLab Rodent Diet 20) and water. Exposure to cold stress was conducted between 8:00 am and 8:30 am for flow cytometry experiments. Pregnant mice were anaesthetized with isoflurane and immediately euthanized by cervical dislocation followed by decapitation to facilitate embryo collection. All mouse embryos were genotyped for sex. For microglial morphological experiments and functional analysis, two embryos of each sex were analyzed across three independent litters for both control and prenatal maternal cold stress groups (n=6 embryos per sex/treatment from N=3 litters). For flow cytometry phagocytosis assays, the entire litter was analyzed.

### DNA extraction and genotyping

Tails were collected from E15.5 mouse embryos and incubated overnight at 56°C in extraction buffer containing: 100 mM NaCl, 50 mM Tris, 100 mM EDTA, and 1% SDS, supplemented with proteinase K (8 units, ∼0.4 mg mL ¹: New England Biolabs P8107S). Genomic DNA was isolated from the digested tissue by precipitation with a saturated NaCl solution (∼6 M), followed by centrifugation at 20,000 × g for 10 min. DNA was subsequently precipitated from the supernatant using isopropanol, and pellets were collected by centrifugation at 20,000 × g for 5 min. Pellets were washed with 70% ethanol and dried at 37°C. Dried DNA was resuspended in 200 µL Tris–EDTA buffer (10 mM Tris, 1 mM EDTA, pH 8) and incubated at 68°C for 1 hr to ensure complete dissolution.

For polymerase chain reaction (PCR) amplification, 1.6 µL of genomic DNA was added to OneTaq® Quick-Load® 2X Master Mix with Standard Buffer (New England Biolabs M0486S), along with 1 µM of primers targeting the sex-linked genes *Sly* and *Xlr* (106). PCR reactions were performed on an Applied Biosystems™ MiniAmp™ Thermal Cycler (Applied Biosystems) using the following conditions: an initial denaturation at 95°C for 3 min, followed by 35 cycles of denaturation at 95°C for 30 sec, annealing at 57°C for 30 sec, and extension at 72°C for 30 sec, with a final extension step at 72°C for 5 min. Amplified PCR products were run on a 1.5% agarose gel prepared in TAE buffer (0.4 M Tris, 0.2 M acetic acid, 10 mM EDTA, pH 8) and visualized using SmartGlow pre-stain for nucleic acid gels (Accuris Instruments E4500-PS) on a Fisherbrand™ Real-Time Electrophoresis System (Fisher Scientific). Fragment sizes were determined using a GeneRuler Ready-to-Use 100 bp DNA Ladder (Thermo Fisher Scientific SM0243).

### Tissue preparation and immunofluorescence staining

E15.5 mouse brains were collected in ice-cold phosphate-buffered saline (PBS) and fixed overnight in 4% paraformaldehyde (PFA) at 4°C. Embryonic brains were then washed in PBS and equilibrated overnight at 4°C in 30% sucrose diluted in PBS. Embryonic brains were embedded in Clear Frozen Section Compound (VWR 95057-838). Serial coronal cryosections (30 μm) were collected through the rostro–caudal axis, spanning the entire hypothalamus, using a Leica CM1950 cryostat (Nussloch, Germany). Coronal sections were serially mounted onto Superfrost Plus glass slides (VWR 48311-703) across five slides (12 sections per slide) for the hypothalamus.

E15.5 hypothalamic cryosections were rehydrated in PBS for 10 min and washed three times for 10 min each in PBS containing 0.1% Triton X-100 (PBST). Cryosections were permeabilized for 30 min in 1X PBS supplemented with 1% Triton X-100, followed by blocking for 1 hr at room temperature (RT) in PBST containing 5% normal donkey serum (NDS, Sigma). Cryosections were incubated overnight at 4°C in PBST with 5% NDS and antibodies against IBA1 (1:200 goat, Invitrogen PA5-18039), AVP (1:400 rabbit, Abcam ab213708), and/or POMC (pro-opiomelanocortin; 1:200 rabbit, Phoenix Pharmaceuticals H-029-30). After incubation, cryosections were washed 4 times for 10 min with PBST and exposed to secondary antibody, including Alexa 488 (Invitrogen anti-goat, A11055) and/or Alexa 594 (Invitrogen anti-rabbit, A21207), diluted in PBST and 5% NDS and left to incubate for 2 hrs at RT. Cryosections were then washed 4 times for 10 min with PBST. Nuclei were stained with Hoechst 33342 (1:1000, Invitrogen) in PBST for 5 min at RT and washed 3 × 5 min with PBST. Cryosections were mounted using Aqua Poly/Mount (Polysciences 18606).

For whole hypothalamic analysis, one slide (12 sections) was analyzed from each embryo. For nuclei specific analysis, one slide was analyzed per embryo, with 2-3 sections assessed for the PVN and 4-5 sections assessed for the ARC based on fluorescent labels. Fluorescent images were acquired with a Zeiss LSM 900 confocal microscope and analyzed using ZEN 3.8 software. Images were exported as RGB (red, green, blue) *.tiffs* for downstream analysis.

### Microglial morphological clustering analysis

Regions of interest (ROIs) were identified in Fiji/ImageJ (v2.16.0, NIH), using the *Polygon Selection* tool to isolate microglia within the ROI (i.e., hypothalamus, PVN or ARC), while excluding tissue outside the target boundaries. For the hypothalamus, ROIs were confined to the hypothalamic area, avoiding adjacent thalamic tissue above the third ventricle. For the PVN, ROIs were restricted to regions containing AVP+ neurons. For the ARC, ROIs were restricted to regions containing POMC+ neurons. Only microglial cells whose soma fell entirely within these marker-defined ROIs were included in downstream analyses to ensure spatial specificity and consistency across sections.

Following methodology adapted from Kim et al. (2024), all ROIs obtained from confocal microscopy images were converted to binary masks using the IBA1 channel in Fiji. Briefly, images were converted to an 8-bit greyscale. Brightness and contrast settings were uniformly applied using the *Enhance Contrast* function, with a saturation threshold set at 0.35%. A global threshold range of 50–255, 68–255, and 80–255 greyscale intensity was applied to the hypothalamus, PVN, and ARC, respectively. Threshold values were optimized for each region to maximize retention of microglial signal while minimizing background fluorescence. Noise reduction was performed using the *Despeckle* function, followed by conversion to a binary mask using the default method (set for dark background images) in Fiji. Remaining small artifacts were removed with the *Remove Outliers* function with default parameters (radius=2 pixels, threshold=50, targeting bright pixels). Binary masks were further refined using the *Analyze Particles* function, which excluded particles below 600 pixels in size to eliminate non-cellular debris and small artifacts. The *Close*–function was applied to remove small holes and artifacts from image processing (79).

Subsequent processing was performed to refine the binary masks by removing particles lacking clearly defined nuclei and separating conjoined cells using a 2-pixel width pencil tool. In cases where the binary did not accurately capture the morphological attributes of the microglia (i.e., less than 70% of the microglia soma and branches were retained), the microglia was removed. This step was blinded, utilizing a custom macro script that iterated through each binary particle, allowing for direct comparison to the corresponding original RGB region and ensuring its completeness and accuracy. The macro script, along with a tutorial video, as well as protocol for inclusion and exclusion of microglia can be found in the ‘Protocol’ section within the ‘Static Microglial Morphology Analysis’ folder of the published GitHub.

Following image processing, microglia were then morphologically clustered following two distinct pipelines utilizing an adapted unsupervised clustering protocol (79), and supervised feature selection clustering protocol (85).

### Unsupervised clustering analysis

The first approach adapted a published pipeline developed by Kim et al. (79). In brief, individual cells were separated, skeletonized, and analyzed using *FracLac* plugin to extract 27 distinct morphological features in Fiji (v2.16.0, NIH). Results from this analysis were then imported into R (v4.4.1, R Core Team, 2024), merged, and compiled into a comprehensive data frame. Dimensionality reduction was performed using Principal Component Analysis (PCA), and K-means clustering was applied using the first two principal components. Since K-means clustering requires the number of clusters to be predefined, the optimal number was determined using the elbow method (within-cluster sum of squares), silhouette analysis, as well as visual inspection of the resulting clusters to ensure they captured biologically distinct groupings. Three distinct morphological clusters were identified. Cluster composition was analyzed as proportions using the *MicrogliaMorphologyR* package in R following the protocol published by Kim et al. (79).

### Supervised clustering analysis

The second approach adapted a published pipeline developed by Maya-Arteaga et al. (85). In brief, individual microglia were separated, skeletonized, and their morphological features quantified using the MorphoGlia App (85), in Python (Python 3.12.2, Python Software Foundation). Feature selection was performed using recursive feature elimination (RFE), where 16 out of 32 features were retained. For this implementation, RFE was supervised using control and cold stress labels as the target variables. Analysis was focused in the PVN for both sexes. Following feature selection, UMAP dimensionality reduction was performed with parameters set with *n* neighbors equal to 30, and minimum distance equal to 0.1. Pseudo-coloring confirmed the unique nature of these clusters. HDBSCAN clustering was then applied with a minimum cluster size of 150 and minimum samples set to 5, resulting in the identification of five distinct clusters.

### Analysis of microglia-AVP neuron interactions

Using the same PVN sections analyzed for morphological clustering, microglia–AVP neuron interactions within the PVN were assessed. Interactions were initially identified in ZEN 3.8 software from 2D 20× immunofluorescence images acquired on a Zeiss LSM 900 confocal microscope. To validate interaction classification, interactions in both sexes from control and cold stress groups were re-examined in ZEN 3.8 software from high-resolution 63× with 3D Z-stacks acquired using a Zeiss LSM 900 confocal microscope. These were further assessed in Imaris (Imaris v10.2, Oxford Instruments) to enable volumetric visualization and confirmation of cell–cell contacts.

Contacts between microglia and AVP+ neurons were classified as touching or wrapping. Contacts were classified as touching if microglia were adjacent to neurons and exhibited limited contact (e.g., a single microglial process contacting the neuronal soma) or were in contact without substantial soma overlap. Contacts were classified as wrapping if there was clear soma-to-soma interactions and/or multiple microglial processes extending and significantly overlapping and wrapping around neuronal structures. Microglial interactions were then mapped back to their original morphological clusters to assess differences in the number and type of interactions between experimental groups and across clusters.

### Analysis of microglial phagocytosis

Using the same PVN sections analyzed for morphological clustering, microglial phagocytic activity within the PVN was assessed. Phagocytic features were evaluated using ZEN 3.8 software in 3D and acquired at 20× with optical zoom using a Zeiss LSM 900 confocal microscope. Microglia were assessed for morphological evidence of phagocytic cups. Evidence of phagocytic cups was identified by the presence of Hoechst+ nuclear material and a circular morphology, often at the distal ends of microglial branches. Microglia were categorized as phagocytic if they possessed one or more clear phagocytic cup(s). Microglia were subsequently mapped back to their original morphological clusters to assess group differences in phagocytosis.

### Flow cytometry analysis of microglial bioparticle engulfment

Pregnant mice were euthanized 3 hrs following their last cold stress exposure on E15.5, and embryos were collected in PBS on ice. Hypothalamic microdissections were performed immediately following euthanasia in PBS on ice and dissected tissues were transferred to individual wells of a 48-well plate containing 270 μL of warmed live-cell culture media supplemented with 30 μL pHrodo™ Deep Red *E. coli* BioParticles™ Conjugate for Phagocytosis (Invitrogen P35360, Thermo Fisher Scientific). Live-cell culture media was prepared by combining: 2.18 mL DMEM (Gibco 11965-092, Thermo Fisher Scientific), 1.09 mL F12 (Gibco 11765-054, Thermo Fisher Scientific), 190 μL Fetal bovine serum (FBS) (Gibco A5670101, Thermo Fisher Scientific), 190 μL horse serum (Gibco 26050-070, Thermo Fisher Scientific), 76 μL B27 supplement (Gibco 17504-044, Thermo Fisher Scientific), 38 μL N2 supplement (Gibco 17502-048, Thermo Fisher Scientific), and 38 μL GlutaMAX (Gibco 35050-061, Thermo Fisher Scientific). pHrodo™ bioparticles were prepared by reconstituting one tube (2 mg) in 2 mL of DMEM (Gibco 11965-092, Thermo Fisher Scientific), followed by sonication, vortexing, and addition to live-cell media.

Hypothalamic tissue was incubated with live-cell media containing bioparticles for 2.5 hrs at 37 °C with 5% CO_2_ in a humidified chamber. After incubation, media containing hypothalamic tissue was mixed by pipetting up and down 40 times within each well to dissociate cells. Cell suspensions were filtered through 35 μm cell strainers (Falcon 352235) into 1.5 mL tubes and centrifuged at 300 × g for 5 min at RT. The supernatant was removed and the resulting pellet was resuspended in 300 μL of HBSS supplemented with 5% FBS (Thermo Fisher Scientific) at RT. The suspension was mixed by pipetting up and down 20 times, filtered through a 35 μm Falcon cell strainer, and transferred into 5 mL round-bottom tubes (VWR 21008-948) prior to flow cytometry analysis. Cell suspensions were analyzed using a Beckman Coulter Life Sciences CytoFLEX LX and Beckman Coulter CytExpert software in the University of British Columbia’s Flow Core Facility (Beckman Coulter Life Sciences; RRID:SCR_017217). Gates were applied uniformly across all experimental samples (Figure S5A–C). The proportion of GFP+ cells expressing allophycocyanin fluorescence (APC+) was quantified as the measure of microglial bioparticle engulfment (Figure S5C).

### Quantification and statistical analysis

All quantifications were visually inspected to ensure that embryos from the independent litters fell within the normal spread of all other data points. For morphological clustering using the unsupervised pipeline, cluster composition was expressed as proportions per embryo, represented as box-and-whisker plots showing median, interquartile range, and minimum-maximum values, and analyzed using a generalized linear mixed-effects model (GLMM) with beta distribution and embryo ID as a random effect in R (version 4.4.1), using the *MicrogliaMorphologyR* package (79). Post hoc comparisons were conducted using Sidak-adjustment. For supervised morphological clustering, group differences in cluster distributions were assessed using a chi-square test, displayed as chi-square residual heatmaps, and analyzed by calculating standardized residuals to characterize the direction and magnitude of associations in R (version 4.4.1).

Microglia–neuron interactions and prevalence of phagocytic microglia from static, single time-point 2D analyses, as well as flow cytometry bioparticle assay quantifications, were represented as bar graphs showing means ± SEM and analyzed using a Student’s t-test or two-way ANOVA (analysis of variance) with Tukey-adjusted post hoc comparisons in GraphPad Prism 9.5.1.

## Declarations

### Ethics approval and consent to participate

Animal work was carried out in accordance with the guidelines and regulations of the Canadian Council on Animal Care and received prior approval from UBC’s Animal Care Committee (protocols A21-0170 and A21-0171). All experimental work was conducted in accordance with an approved Biosafety Protocol (B21-0133).

### Consent for publication

Not applicable.

### Availability of data and materials

Original images used for analyses across the current study are available upon request to the lead contact. The datasets generated and/or analyzed during the current study, alongside a video tutorial, are available in the GitHub repository: https://github.com/RosinLabUBC/Microglia-Cold-Stress-Project-AL

### Competing interests

The authors declare that they have no competing interests.

### Funding

Alexandra Lawson was supported by the Canadian Neurodevelopmental Research Training Platform (CanNRT). This work was supported by a Canadian Institutes of Health Research (CIHR) Project Grant (GR028789) to Jessica M. Rosin. Jessica M. Rosin is a Michael Smith Health Research (MSHR) BC Scholar and a Tier 2 Canada Research Chair (CRC) in Immune Regulation of Developmental Programs.

### Authors’ contributions

JMR conceived and designed the study, assisted with hypothalamic microdissections, and advised on all data analysis approaches. MR assisted with mouse handling, tissue sectioning, immunofluorescence staining, flow cytometry, and microscopy. AL conducted mouse handling, flow cytometry acquisition and analysis, microscopy and associated analyses, and adapted morphologic analysis code for embryonic microglia, in addition to performing all data analysis and statistical assessment. AL and JMR drafted the original manuscript, with edits incorporated from AL, MR, and JMR. All authors read and approved the final manuscript.

## Acknowledgements

The authors thank the University of British Columbia (UBC) Centre for Disease Modelling for animal care, Quincy Collins and other members of the Rosin laboratory for help with immunofluorescence staining, Erich Giedzinki from *Imaris* as well as Guang Gao from the LSI Imaging core for technical guidance with *Imaris,* Andrew Johnson from the UBC Flow Core Facility for technical support during flow cytometry experiments, and Biljana Stojkova from the Applied Statistics and Data Science group for statistical guidance. We also thank Annie Ciernia and Jennifer Kim for critical discussions and guidance. Figures 1A and 6K were created with BioRender.com. Portions of this manuscript appear in AL’s Master of Science thesis at UBC.

## Abbreviations

2D / 3D: two-/three-dimensional
ADHD: attention-deficit/hyperactivity disorder
ANOVA: analysis of variance
APC: allophycocyanin
ARC: arcuate nucleus
ASD: autism spectrum disorder
AVP: arginine vasopressin
bp: base pair
BPA: bisphenol A
CO2: carbon dioxide
CRH: corticotropin-releasing hormone
Cx3cr1: CX3C chemokine receptor 1 (gene/locus)
DMEM: Dulbecco’s Modified Eagle Medium
DNA: deoxyribonucleic acid
E (e.g., E11.5, E15.5): embryonic day
EDTA: ethylenediaminetetraacetic acid
eGFP/GFP: (enhanced) green fluorescent protein
FBS: fetal bovine serum
GLMM: generalized linear mixed-effects model
HBSS: Hanks’ Balanced Salt Solution
HDBSCAN: Hierarchical Density-Based Spatial Clustering of Applications with Noise
HYPO: hypothalamus
IBA1: ionized calcium-binding adapter molecule 1
LSI: Life Sciences Institute
MSHR: Michael Smith Health Research (BC)
NaCl: sodium chloride
NDDs: neurodevelopmental disorders
NDS: normal donkey serum
NIH: National Institutes of Health
OXT: oxytocin
PBS: phosphate-buffered saline
PBST: PBS + Triton X-100
PC(s): principal component(s)
PCA: principal component analysis
PCR: polymerase chain reaction
PFA: paraformaldehyde
pHrodo: pH-sensitive bioparticle conjugate (trade name)
POMC: pro-opiomelanocortin
PTSD: post-traumatic stress disorder
RFE: recursive feature elimination
RGB: red, green, blue
ROIs: regions of interest
RRID: Research Resource Identifier
RT: room temperature
SDS: sodium dodecyl sulfate
SEM: standard error of the mean
Sly: sex-linked genotyping gene
TAE: Tris-acetate-EDTA (buffer)
Tris: tris(hydroxymethyl)aminomethane
UBC: University of British Columbia
UMAP: Uniform Manifold Approximation and Projection
WT: wild-type
χ²: chi-square

## Notes

### Competing Interest Statement

The authors have declared no competing interest.

### Summary of Updates

All aspects of the manuscript (text, figures, supplemental materials) have been revised.

