## Supplementary Tables 1-19 for "Fetal microglia show region-specific and morphology-dependent sex differences in their responsiveness to prenatal maternal stress"

**Supplementary Table 1. Two-way ANOVA results for the number of microglial cells per mouse in the E15.5 hypothalamus**

| Number of Microglial Cells in the Hypothalamus | | |
|---|---|---|
| **Tukey-adjusted post hoc comparisons** | **Adjusted p value** | **Summary** |
| Male: Control vs. Male: Cold Stress | 0.9803 | ns |
| Male: Control vs. Female: Control | 0.8394 | ns |
| Male: Control vs. Female: Cold Stress | 0.8951 | ns |
| Male: Cold Stress vs. Female: Control | 0.6255 | ns |
| Male: Cold Stress vs. Female: Cold Stress | 0.7021 | ns |
| Female: Control vs. Female: Cold Stress | 0.9992 | ns |
| **Two-way ANOVA** | **p value** | **Summary** |
| Treatment | 0.7212 | ns |
| Sex | 0.1906 | ns |
| Interaction | 0.8587 | ns |

**Supplementary Table 2. GLMM results for microglial cluster proportions across rostro-caudal hypothalamic sections in control embryos**

| Section 01 | | | | |
|---|---|---|---|---|
| **Sidak-adjusted post hoc comparisons** | | | **Adjusted p value** | **Summary** |
| Ameboid: Male vs. Female | | | 0.2254 | ns |
| Ramified: Male vs. Female | | | 0.9998 | ns |
| Rod-Like: Male vs. Female | | | 0.1819 | ns |
| **Type II Wald Chi-square tests** | **Chi-sq** | **Df** | **p value** | **Summary** |
| Sex | 0.0027 | 1 | 0.9588 | ns |
| Cluster | 13.35 | 2 | 0.0013 | ** |
| Interaction | 6.445 | 2 | 0.0399 | * |
| **Section 02** | | | | |
| **Sidak-adjusted post hoc comparisons** | | | **Adjusted p value** | **Summary** |
| Ameboid: Male vs. Female | | | 0.2602 | ns |
| Ramified: Male vs. Female | | | 0.7950 | ns |
| Rod-Like: Male vs. Female | | | 0.8500 | ns |
| **Type II Wald Chi-square tests** | **Chi-sq** | **Df** | **p value** | **Summary** |
| Sex | 0.0052 | 1 | 0.9428 | ns |
| Cluster | 41.40 | 2 | <0.0001 | **** |
| Interaction | 3.976 | 2 | 0.1370 | ns |
| **Section 03** | | | | |
| **Sidak-adjusted post hoc comparisons** | | | **Adjusted p value** | **Summary** |
| Ameboid: Male vs. Female | | | 0.6133 | ns |
| Ramified: Male vs. Female | | | 0.8006 | ns |
| Rod-Like: Male vs. Female | | | >0.999 | ns |
| **Type II Wald Chi-square tests** | **Chi-sq** | **Df** | **p value** | **Summary** |
| Sex | 0.0182 | 1 | 0.8927 | ns |
| Cluster | 12.244 | 2 | 0.0022 | ** |
| Interaction | 1.8552 | 2 | 0.3955 | ns |
| **Section 04** | | | | |
| **Sidak-adjusted post hoc comparisons** | | | **Adjusted p value** | **Summary** |
| Ameboid: Male vs. Female | | | 0.9811 | ns |
| Ramified: Male vs. Female | | | 0.9867 | ns |
| Rod-Like: Male vs. Female | | | 0.8194 | ns |
| **Type II Wald Chi-square tests** | **Chi-sq** | **Df** | **p value** | **Summary** |
| Sex | 0.0012 | 1 | 0.9722 | ns |
| Cluster | 41.739 | 2 | <0.0001 | **** |
| Interaction | 0.8158 | 2 | 0.6650 | ns |
| **Section 05** | | | | |
| **Sidak-adjusted post hoc comparisons** | | | **Adjusted p value** | **Summary** |
| Ameboid: Male vs. Female | | | 0.8639 | ns |
| Ramified: Male vs. Female | | | 0.9916 | ns |
| Rod-Like: Male vs. Female | | | 0.9616 | ns |
| **Type II Wald Chi-square tests** | **Chi-sq** | **Df** | **p value** | **Summary** |

| | | | | |
|---|---|---|---|---|
| Sex | <0.0001 | 1 | 0.9986 | ns |
| Cluster | 48.371 | 2 | <0.0001 | **** |
| Interaction | 0.7429 | 2 | 0.6897 | ns |
| **Section 06** | | | | |
| **Sidak-adjusted post hoc comparisons** | | | **Adjusted p value** | **Summary** |
| Ameboid: Male vs. Female | | | 0.6512 | ns |
| Ramified: Male vs. Female | | | 0.8171 | ns |
| Rod-Like: Male vs. Female | | | 0.9877 | ns |
| **Type II Wald Chi-square tests** | **Chi-sq** | **Df** | **p value** | **Summary** |
| Sex | 0.0016 | 1 | 0.9682 | ns |
| Cluster | 36.263 | 2 | <0.0001 | **** |
| Interaction | 1.7926 | 2 | 0.4081 | ns |
| **Section 07** | | | | |
| **Sidak-adjusted post hoc comparisons** | | | **Adjusted p value** | **Summary** |
| Ameboid: Male vs. Female | | | 0.9049 | ns |
| Ramified: Male vs. Female | | | 0.9275 | ns |
| Rod-Like: Male vs. Female | | | 0.9999 | ns |
| **Type II Wald Chi-square tests** | **Chi-sq** | **Df** | **p value** | **Summary** |
| Sex | 0.004 | 1 | 0.9494 | ns |
| Cluster | 14.044 | 2 | 0.0009 | *** |
| Interaction | 0.6703 | 2 | 0.7152 | ns |
| **Section 08** | | | | |
| **Sidak-adjusted post hoc comparisons** | | | **Adjusted p value** | **Summary** |
| Ameboid: Male vs. Female | | | 0.9995 | ns |
| Ramified: Male vs. Female | | | 1.0000 | ns |
| Rod-Like: Male vs. Female | | | 1.0000 | ns |
| **Type II Wald Chi-square tests** | **Chi-sq** | **Df** | **p value** | **Summary** |
| Sex | 0.0026 | 1 | 0.9595 | ns |
| Cluster | 26.569 | 2 | <0.0001 | **** |
| Interaction | 0.0010 | 2 | 0.9947 | ns |
| **Section 09** | | | | |
| **Sidak-adjusted post hoc comparisons** | | | **Adjusted p value** | **Summary** |
| Ameboid: Male vs. Female | | | 0.7973 | ns |
| Ramified: Male vs. Female | | | 0.9952 | ns |
| Rod-Like: Male vs. Female | | | 0.9327 | ns |
| **Type II Wald Chi-square tests** | **Chi-sq** | **Df** | **p value** | **Summary** |
| Sex | 0.0022 | 1 | 0.9623 | ns |
| Cluster | 10.785 | 2 | 0.0046 | ** |
| Interaction | 0.9996 | 2 | 0.6067 | ns |
| **Section 10** | | | | |
| **Sidak-adjusted post hoc comparisons** | | | **Adjusted p value** | **Summary** |
| Ameboid: Male vs. Female | | | 0.9992 | ns |
| Ramified: Male vs. Female | | | 0.9834 | ns |

| | | | | |
|---|---|---|---|---|
| Rod-Like: Male vs. Female | | | 0.9836 | ns |
| **Type II Wald Chi-square tests** | **Chi-sq** | **Df** | **p value** | **Summary** |
| Sex | 0.0072 | 1 | 0.9326 | ns |
| Cluster | 28.181 | 2 | <0.0001 | **** |
| Interaction | 0.2169 | 2 | 0.8972 | ns |
| **Section 11** | | | | |
| **Sidak-adjusted post hoc comparisons** | | | **Adjusted p value** | **Summary** |
| Ameboid: Male vs. Female | | | 0.9723 | ns |
| Ramified: Male vs. Female | | | 0.8907 | ns |
| Rod-Like: Male vs. Female | | | 1.0000 | ns |
| **Type II Wald Chi-square tests** | **Chi-sq** | **Df** | **p value** | **Summary** |
| Sex | 0.0030 | 1 | 0.8606 | ns |
| Cluster | 14.433 | 2 | 0.0007 | *** |
| Interaction | 0.5308 | 2 | 0.7669 | ns |
| **Section 12** | | | | |
| **Sidak-adjusted post hoc comparisons** | | | **Adjusted p value** | **Summary** |
| Ameboid: Male vs. Female | | | 0.9683 | ns |
| Ramified: Male vs. Female | | | 0.9778 | ns |
| Rod-Like: Male vs. Female | | | 0.8688 | ns |
| **Type II Wald Chi-square tests** | **Chi-sq** | **Df** | **p value** | **Summary** |
| Sex | 0.0027 | 1 | 0.9586 | ns |
| Cluster | 53.581 | 2 | <0.0001 | **** |
| Interaction | 0.7655 | 2 | 0.6820 | ns |

**Supplementary Table 3. GLMM results for microglial cluster proportions across 12 rostro-caudal hypothalamic sections in male embryos**

| | | | | |
|---|---|---|---|---|
| **Section 01 (males)** | | | | |
| **Sidak-adjusted post hoc comparisons** | | | **Adjusted p value** | **Summary** |
| Ameboid: Control vs. Cold Stress | | | 0.9979 | ns |
| Ramified: Control vs. Cold Stress | | | 0.8644 | ns |
| Rod-Like: Control vs. Cold Stress | | | 0.7786 | ns |
| **Type II Wald Chi-square tests** | **Chi-sq** | **Df** | **p value** | **Summary** |
| Treatment | 0.0019 | 1 | 0.9651 | ns |
| Cluster | 12.5486 | 2 | 0.0019 | ** |
| Interaction | 1.2323 | 2 | 0.54 | ns |
| **Section 02 (males)** | | | | |
| **Sidak-adjusted post hoc comparisons** | | | **Adjusted p value** | **Summary** |
| Ameboid: Control vs. Cold Stress | | | 0.9968 | ns |
| Ramified: Control vs. Cold Stress | | | 0.7688 | ns |
| Rod-Like: Control vs. Cold Stress | | | 0.8883 | ns |
| **Type II Wald Chi-square tests** | **Chi-sq** | **Df** | **p value** | **Summary** |
| Treatment | 0.0037 | 1 | 0.9517 | ns |
| Cluster | 22.9026 | 2 | <0.0001 | **** |
| Interaction | 1.1981 | 2 | 0.5493 | ns |
| **Section 03 (males)** | | | | |
| **Sidak-adjusted post hoc comparisons** | | | **Adjusted p value** | **Summary** |
| Ameboid: Control vs. Cold Stress | | | 0.5995 | ns |
| Ramified: Control vs. Cold Stress | | | 0.6801 | ns |
| Rod-Like: Control vs. Cold Stress | | | >0.9999 | ns |
| **Type II Wald Chi-square tests** | **Chi-sq** | **Df** | **p value** | **Summary** |
| Treatment | 0.0014 | 1 | 0.9704 | ns |
| Cluster | 8.1666 | 2 | 0.0169 | * |
| Interaction | 2.2591 | 2 | 0.3232 | ns |
| **Section 04 (males)** | | | | |
| **Sidak-adjusted post hoc comparisons** | | | **Adjusted p value** | **Summary** |
| Ameboid: Control vs. Cold Stress | | | 0.8133 | ns |
| Ramified: Control vs. Cold Stress | | | 0.4200 | ns |
| Rod-Like: Control vs. Cold Stress | | | 0.7294 | ns |
| **Type II Wald Chi-square tests** | **Chi-sq** | **Df** | **p value** | **Summary** |
| Treatment | 0.0127 | 1 | 0.9102 | ns |
| Cluster | 40.9989 | 2 | <0.0001 | **** |
| Interaction | 3.3943 | 2 | 0.1832 | ns |
| **Section 05 (males)** | | | | |
| **Sidak-adjusted post hoc comparisons** | | | **Adjusted p value** | **Summary** |
| Ameboid: Control vs. Cold Stress | | | 0.9673 | ns |
| Ramified: Control vs. Cold Stress | | | 0.8933 | ns |

| | | | | |
|---|---|---|---|---|
| Rod-Like: Control vs. Cold Stress | | | 0.9979 | ns |
| **Type II Wald Chi-square tests** | **Chi-sq** | **Df** | **p value** | **Summary** |
| Treatment | 0.0038 | 1 | 0.9511 | ns |
| Cluster | 29.7314 | 2 | <0.0001 | **** |
| Interaction | 0.5948 | 2 | 0.7427 | ns |
| **Section 06 (males)** | | | | |
| **Sidak-adjusted post hoc comparisons** | | | **Adjusted p value** | **Summary** |
| Ameboid: Control vs. Cold Stress | | | >0.9999 | ns |
| Ramified: Control vs. Cold Stress | | | >0.9999 | ns |
| Rod-Like: Control vs. Cold Stress | | | 0.9922 | ns |
| **Type II Wald Chi-square tests** | **Chi-sq** | **Df** | **p value** | **Summary** |
| Treatment | 0.025 | 1 | 0.8744 | ns |
| Cluster | 29.2279 | 2 | <0.0001 | **** |
| Interaction | 0.0397 | 2 | 0.9804 | ns |
| **Section 07 (males)** | | | | |
| **Sidak-adjusted post hoc comparisons** | | | **Adjusted p value** | **Summary** |
| Ameboid: Control vs. Cold Stress | | | 0.9903 | ns |
| Ramified: Control vs. Cold Stress | | | 0.9980 | ns |
| Rod-Like: Control vs. Cold Stress | | | 0.9763 | ns |
| **Type II Wald Chi-square tests** | **Chi-sq** | **Df** | **p value** | **Summary** |
| Treatment | 0.0033 | 1 | 0.9545 | ns |
| Cluster | 18.6004 | 2 | <0.0001 | **** |
| Interaction | 0.2306 | 2 | 0.8911 | ns |
| **Section 08 (males)** | | | | |
| **Sidak-adjusted post hoc comparisons** | | | **Adjusted p value** | **Summary** |
| Ameboid: Control vs. Cold Stress | | | 0.7863 | ns |
| Ramified: Control vs. Cold Stress | | | 0.9880 | ns |
| Rod-Like: Control vs. Cold Stress | | | 0.9428 | ns |
| **Type II Wald Chi-square tests** | **Chi-sq** | **Df** | **p value** | **Summary** |
| Treatment | 0.0011 | 1 | 0.973 | ns |
| Cluster | 16.7793 | 2 | 0.0002 | *** |
| Interaction | 1.0389 | 2 | 0.5949 | ns |
| **Section 09 (males)** | | | | |
| **Sidak-adjusted post hoc comparisons** | | | **Adjusted p value** | **Summary** |
| Ameboid: Control vs. Cold Stress | | | 0.9158 | ns |
| Ramified: Control vs. Cold Stress | | | 0.9819 | ns |
| Rod-Like: Control vs. Cold Stress | | | 0.6936 | ns |
| **Type II Wald Chi-square tests** | **Chi-sq** | **Df** | **p value** | **Summary** |
| Treatment | 0.0001 | 1 | 0.9928 | ns |
| Cluster | 14.1057 | 2 | 0.0009 | *** |
| Interaction | 1.4145 | 2 | 0.493 | ns |

| | | | | |
|---|---|---|---|---|
| **Section 10 (males)** | | | | |
| **Sidak-adjusted post hoc comparisons** | | | **Adjusted p value** | **Summary** |
| Ameboid: Control vs. Cold Stress | | | 0.7886 | ns |
| Ramified: Control vs. Cold Stress | | | 0.9782 | ns |
| Rod-Like: Control vs. Cold Stress | | | 0.9347 | ns |
| **Type II Wald Chi-square tests** | **Chi-sq** | **Df** | **p value** | **Summary** |
| Treatment | 0.0019 | 1 | 0.965 | ns |
| Cluster | 24.6605 | 2 | <0.0001 | **** |
| Interaction | 1.1004 | 2 | 0.5768 | ns |
| **Section 11 (males)** | | | | |
| **Sidak-adjusted post hoc comparisons** | | | **Adjusted p value** | **Summary** |
| Ameboid: Control vs. Cold Stress | | | 0.7665 | ns |
| Ramified: Control vs. Cold Stress | | | 0.6079 | ns |
| Rod-Like: Control vs. Cold Stress | | | 0.9995 | ns |
| **Type II Wald Chi-square tests** | **Chi-sq** | **Df** | **p value** | **Summary** |
| Treatment | 0.0158 | 1 | 0.8999 | ns |
| Cluster | 17.6191 | 2 | 0.0001 | *** |
| Interaction | 1.9781 | 2 | 0.3719 | ns |
| **Section 12 (males)** | | | | |
| **Sidak-adjusted post hoc comparisons** | | | **Adjusted p value** | **Summary** |
| Ameboid: Control vs. Cold Stress | | | 0.9400 | ns |
| Ramified: Control vs. Cold Stress | | | 0.8046 | ns |
| Rod-Like: Control vs. Cold Stress | | | 0.9999 | ns |
| **Type II Wald Chi-square tests** | **Chi-sq** | **Df** | **p value** | **Summary** |
| Treatment | 0.0286 | 1 | 0.8657 | ns |
| Cluster | 16.5569 | 2 | 0.0003 | *** |
| Interaction | 0.8888 | 2 | 0.6412 | ns |

**Supplementary Table 4. GLMM results for microglial cluster proportions across 12 rostro-caudal hypothalamic sections in female embryos**

| | | | | |
|---|---|---|---|---|
| **Section 01 (females)** | | | | |
| **Sidak-adjusted post hoc comparisons** | | | **Adjusted p value** | **Summary** |
| Ameboid: Control vs. Cold Stress | | | 0.8439 | ns |
| Ramified: Control vs. Cold Stress | | | 0.6490 | ns |
| Rod-Like: Control vs. Cold Stress | | | 0.1556 | ns |
| **Type II Wald Chi-square tests** | **Chi-sq** | **Df** | **p value** | **Summary** |
| Treatment | 0.0001 | 1 | 0.9929 | ns |
| Cluster | 24.541 | 2 | <0.0001 | **** |
| Interaction | 5.328 | 2 | 0.0697 | ns |
| **Section 02 (females)** | | | | |
| **Sidak-adjusted post hoc comparisons** | | | **Adjusted p value** | **Summary** |
| Ameboid: Control vs. Cold Stress | | | 0.9149 | ns |
| Ramified: Control vs. Cold Stress | | | 0.9716 | ns |
| Rod-Like: Control vs. Cold Stress | | | 0.7137 | ns |
| **Type II Wald Chi-square tests** | **Chi-sq** | **Df** | **p value** | **Summary** |
| Treatment | 0.0038 | 1 | 0.9507 | ns |
| Cluster | 30.0313 | 2 | <0.0001 | **** |
| Interaction | 1.3964 | 2 | 0.4975 | ns |
| **Section 03 (females)** | | | | |
| **Sidak-adjusted post hoc comparisons** | | | **Adjusted p value** | **Summary** |
| Ameboid: Control vs. Cold Stress | | | 0.0165 | * |
| Ramified: Control vs. Cold Stress | | | 0.0126 | * |
| Rod-Like: Control vs. Cold Stress | | | 0.9882 | ns |
| **Type II Wald Chi-square tests** | **Chi-sq** | **Df** | **p value** | **Summary** |
| Treatment | 0.0009 | 1 | 0.9761 | ns |
| Cluster | 37.1602 | 2 | <0.0001 | **** |
| Interaction | 15.9706 | 2 | 0.0003 | *** |
| **Section 04 (females)** | | | | |
| **Sidak-adjusted post hoc comparisons** | | | **Adjusted p value** | **Summary** |
| Ameboid: Control vs. Cold Stress | | | 0.3016 | ns |
| Ramified: Control vs. Cold Stress | | | 0.4384 | ns |
| Rod-Like: Control vs. Cold Stress | | | 0.9998 | ns |
| **Type II Wald Chi-square tests** | **Chi-sq** | **Df** | **p value** | **Summary** |
| Treatment | 0 | 1 | 0.9985 | ns |
| Cluster | 63.9773 | 2 | <0.0001 | **** |
| Interaction | 4.3601 | 2 | 0.113 | ns |
| **Section 05 (females)** | | | | |
| **Sidak-adjusted post hoc comparisons** | | | **Adjusted p value** | **Summary** |
| Ameboid: Control vs. Cold Stress | | | 0.8546 | ns |
| Ramified: Control vs. Cold Stress | | | 0.9998 | ns |

| | | | | |
|---|---|---|---|---|
| Rod-Like: Control vs. Cold Stress | | | 0.8604 | ns |
| **Type II Wald Chi-square tests** | **Chi-sq** | **Df** | **p value** | **Summary** |
| Treatment | 0.0017 | 1 | 0.9674 | ns |
| Cluster | 59.5309 | 2 | <0.0001 | **** |
| Interaction | 1.0123 | 2 | 0.6028 | ns |
| **Section 06 (females)** | | | | |
| **Sidak-adjusted post hoc comparisons** | | | **Adjusted p value** | **Summary** |
| Ameboid: Control vs. Cold Stress | | | 0.1549 | ns |
| Ramified: Control vs. Cold Stress | | | 0.6039 | ns |
| Rod-Like: Control vs. Cold Stress | | | 0.8737 | ns |
| **Type II Wald Chi-square tests** | **Chi-sq** | **Df** | **p value** | **Summary** |
| Treatment | 0.0008 | 1 | 0.9777 | ns |
| Cluster | 31.6776 | 2 | <0.0001 | **** |
| Interaction | 5.3927 | 2 | 0.0675 | ns |
| **Section 07 (females)** | | | | |
| **Sidak-adjusted post hoc comparisons** | | | **Adjusted p value** | **Summary** |
| Ameboid: Control vs. Cold Stress | | | 0.3363 | ns |
| Ramified: Control vs. Cold Stress | | | 0.9197 | ns |
| Rod-Like: Control vs. Cold Stress | | | 0.7534 | ns |
| **Type II Wald Chi-square tests** | **Chi-sq** | **Df** | **p value** | **Summary** |
| Treatment | 0.0022 | 1 | 0.9624 | ns |
| Cluster | 10.2485 | 2 | 0.0060 | ** |
| Interaction | 3.4365 | 2 | 0.1794 | ns |
| **Section 08 (females)** | | | | |
| **Sidak-adjusted post hoc comparisons** | | | **Adjusted p value** | **Summary** |
| Ameboid: Control vs. Cold Stress | | | 0.5094 | ns |
| Ramified: Control vs. Cold Stress | | | 0.9639 | ns |
| Rod-Like: Control vs. Cold Stress | | | 0.9146 | ns |
| **Type II Wald Chi-square tests** | **Chi-sq** | **Df** | **p value** | **Summary** |
| Treatment | 0.0207 | 1 | 0.8857 | ns |
| Cluster | 17.7524 | 2 | 0.0001 | *** |
| Interaction | 2.0641 | 2 | 0.3563 | ns |
| **Section 09 (females)** | | | | |
| **Sidak-adjusted post hoc comparisons** | | | **Adjusted p value** | **Summary** |
| Ameboid: Control vs. Cold Stress | | | 0.5936 | ns |
| Ramified: Control vs. Cold Stress | | | 0.9798 | ns |
| Rod-Like: Control vs. Cold Stress | | | 0.8554 | ns |
| **Type II Wald Chi-square tests** | **Chi-sq** | **Df** | **p value** | **Summary** |
| Treatment | 0.0016 | 1 | 0.9678 | ns |
| Cluster | 12.6479 | 2 | 0.0018 | ** |
| Interaction | 1.902 | 2 | 0.3863 | ns |

| | | | | |
|---|---|---|---|---|
| **Section 10 (females)** | | | | |
| **Sidak-adjusted post hoc comparisons** | | | **Adjusted p value** | **Summary** |
| Ameboid: Control vs. Cold Stress | | | 0.8819 | ns |
| Ramified: Control vs. Cold Stress | | | 0.8065 | ns |
| Rod-Like: Control vs. Cold Stress | | | 0.9981 | ns |
| **Type II Wald Chi-square tests** | **Chi-sq** | **Df** | **p value** | **Summary** |
| Treatment | 0.0004 | 1 | 0.985 | ns |
| Cluster | 33.5252 | 2 | <0.0001 | **** |
| Interaction | 1.1052 | 2 | 0.5755 | ns |
| **Section 11 (females)** | | | | |
| **Sidak-adjusted post hoc comparisons** | | | **Adjusted p value** | **Summary** |
| Ameboid: Control vs. Cold Stress | | | 0.8822 | ns |
| Ramified: Control vs. Cold Stress | | | 0.9297 | ns |
| Rod-Like: Control vs. Cold Stress | | | >0.9999 | ns |
| **Type II Wald Chi-square tests** | **Chi-sq** | **Df** | **p value** | **Summary** |
| Treatment | 0.0045 | 1 | 0.9467 | ns |
| Cluster | 39.364 | 2 | <0.0001 | **** |
| Interaction | 0.7279 | 2 | 0.6949 | ns |
| **Section 12 (females)** | | | | |
| **Sidak-adjusted post hoc comparisons** | | | **Adjusted p value** | **Summary** |
| Ameboid: Control vs. Cold Stress | | | 0.9994 | ns |
| Ramified: Control vs. Cold Stress | | | 0.9513 | ns |
| Rod-Like: Control vs. Cold Stress | | | 0.9800 | ns |
| **Type II Wald Chi-square tests** | **Chi-sq** | **Df** | **p value** | **Summary** |
| Treatment | 0.0227 | 1 | 0.8803 | ns |
| Cluster | 37.0275 | 2 | <0.0001 | **** |
| Interaction | 0.3343 | 2 | 0.8461 | ns |

**Supplementary Table 5. GLMM results with section in omnibus for microglial cluster proportions across 12 rostro-caudal hypothalamic sections in control embryos**

| Control Microglial Cluster Proportions (section in omnibus) | | |
|---|---|---|
| **Sidak-adjusted post hoc comparisons** | **Adjusted p value** | **Summary** |
| **Section 01** | | |
| Ameboid: Male vs. Female | 0.9568 | ns |
| Ramified: Male vs. Female | >0.9999 | ns |
| Rod-Like: Male vs. Female | 0.9152 | ns |
| **Section 02** | | |
| Ameboid: Male vs. Female | 0.9951 | ns |
| Ramified: Male vs. Female | >0.9999 | ns |
| Rod-Like: Male vs. Female | >0.9999 | ns |
| **Section 03** | | |
| Ameboid: Male vs. Female | >0.9999 | ns |
| Ramified: Male vs. Female | >0.9999 | ns |
| Rod-Like: Male vs. Female | >0.9999 | ns |
| **Section 04** | | |
| Ameboid: Male vs. Female | >0.9999 | ns |
| Ramified: Male vs. Female | >0.9999 | ns |
| Rod-Like: Male vs. Female | >0.9999 | ns |
| **Section 05** | | |
| Ameboid: Male vs. Female | >0.9999 | ns |
| Ramified: Male vs. Female | >0.9999 | ns |
| Rod-Like: Male vs. Female | >0.9999 | ns |
| **Section 06** | | |
| Ameboid: Male vs. Female | >0.9999 | ns |
| Ramified: Male vs. Female | >0.9999 | ns |
| Rod-Like: Male vs. Female | >0.9999 | ns |
| **Section 07** | | |
| Ameboid: Male vs. Female | >0.9999 | ns |
| Ramified: Male vs. Female | >0.9999 | ns |
| Rod-Like: Male vs. Female | >0.9999 | ns |
| **Section 08** | | |
| Ameboid: Male vs. Female | >0.9999 | ns |
| Ramified: Male vs. Female | >0.9999 | ns |
| Rod-Like: Male vs. Female | >0.9999 | ns |
| **Section 09** | | |
| Ameboid: Male vs. Female | >0.9999 | ns |
| Ramified: Male vs. Female | >0.9999 | ns |
| Rod-Like: Male vs. Female | >0.9999 | ns |
| **Section 10** | | |
| Ameboid: Male vs. Female | >0.9999 | ns |
| Ramified: Male vs. Female | >0.9999 | ns |

| | | |
|---|---|---|
| Rod-Like: Male vs. Female | >0.9999 | ns |
| **Section 11** | | |
| Ameboid: Male vs. Female | >0.9999 | ns |
| Ramified: Male vs. Female | >0.9999 | ns |
| Rod-Like: Male vs. Female | >0.9999 | ns |
| **Section 12** | | |
| Ameboid: Male vs. Female | >0.9999 | ns |
| Ramified: Male vs. Female | >0.9999 | ns |
| Rod-Like: Male vs. Female | >0.9999 | ns |

| **Type II Wald Chi-square tests** | **Chi-sq** | **Df** | **p value** | **Summary** |
|---|---|---|---|---|
| Cluster | 271.3191 | 2 | <0.0001 | **** |
| Sex | 0.0451 | 1 | 0.8317 | ns |
| Section | 0.0612 | 11 | >0.9999 | ns |
| Cluster × Sex | 0.7625 | 2 | 0.6830 | ns |
| Cluster × Section | 51.1091 | 22 | 0.0004 | *** |
| Sex × Section | 0.0319 | 11 | >0.9999 | ns |
| Cluster × Sex × Section | 16.9585 | 22 | 0.7657 | ns |

**Supplementary Table 6. GLMM with section in omnibus results for microglial cluster proportions across 12 rostro-caudal hypothalamic sections in male embryos**

| Male Microglial Cluster Proportions (section in omnibus) | | |
|---|---|---|
| **Sidak-adjusted post hoc comparisons** | **Adjusted p value** | **Summary** |
| **Section 01 (males)** | | |
| Ameboid: Control vs. Cold Stress | >0.9999 | ns |
| Ramified: Control vs. Cold Stress | >0.9999 | ns |
| Rod-Like: Control vs. Cold Stress | >0.9999 | ns |
| **Section 02 (males)** | | |
| Ameboid: Control vs. Cold Stress | >0.9999 | ns |
| Ramified: Control vs. Cold Stress | >0.9999 | ns |
| Rod-Like: Control vs. Cold Stress | >0.9999 | ns |
| **Section 03 (males)** | | |
| Ameboid: Control vs. Cold Stress | >0.9999 | ns |
| Ramified: Control vs. Cold Stress | >0.9999 | ns |
| Rod-Like: Control vs. Cold Stress | >0.9999 | ns |
| **Section 04 (males)** | | |
| Ameboid: Control vs. Cold Stress | >0.9999 | ns |
| Ramified: Control vs. Cold Stress | 0.9998 | ns |
| Rod-Like: Control vs. Cold Stress | >0.9999 | ns |
| **Section 05 (males)** | | |
| Ameboid: Control vs. Cold Stress | >0.9999 | ns |
| Ramified: Control vs. Cold Stress | >0.9999 | ns |
| Rod-Like: Control vs. Cold Stress | >0.9999 | ns |
| **Section 06 (males)** | | |
| Ameboid: Control vs. Cold Stress | >0.9999 | ns |
| Ramified: Control vs. Cold Stress | >0.9999 | ns |
| Rod-Like: Control vs. Cold Stress | >0.9999 | ns |
| **Section 07 (males)** | | |
| Ameboid: Control vs. Cold Stress | >0.9999 | ns |
| Ramified: Control vs. Cold Stress | >0.9999 | ns |
| Rod-Like: Control vs. Cold Stress | >0.9999 | ns |
| **Section 08 (males)** | | |
| Ameboid: Control vs. Cold Stress | >0.9999 | ns |
| Ramified: Control vs. Cold Stress | >0.9999 | ns |
| Rod-Like: Control vs. Cold Stress | >0.9999 | ns |
| **Section 09 (males)** | | |
| Ameboid: Control vs. Cold Stress | >0.9999 | ns |
| Ramified: Control vs. Cold Stress | >0.9999 | ns |
| Rod-Like: Control vs. Cold Stress | >0.9999 | ns |
| **Section 10 (males)** | | |
| Ameboid: Control vs. Cold Stress | >0.9999 | ns |
| Ramified: Control vs. Cold Stress | >0.9999 | ns |

| | | | | |
|---|---|---|---|---|
| Rod-Like: Control vs. Cold Stress | | | >0.9999 | ns |
| **Section 11 (males)** | | | | |
| Ameboid: Control vs. Cold Stress | | | >0.9999 | ns |
| Ramified: Control vs. Cold Stress | | | >0.9999 | ns |
| Rod-Like: Control vs. Cold Stress | | | >0.9999 | ns |
| **Section 12 (males)** | | | | |
| Ameboid: Control vs. Cold Stress | | | >0.9999 | ns |
| Ramified: Control vs. Cold Stress | | | >0.9999 | ns |
| Rod-Like: Control vs. Cold Stress | | | >0.9999 | ns |
| **Type II Wald Chi-square tests** | **Chi-sq** | **Df** | **p value** | **Summary** |
| Cluster | 201.2410 | 2 | <0.0001 | **** |
| Treatment | 0.0018 | 1 | 0.9658 | ns |
| Section | 0.0342 | 11 | >0.9999 | ns |
| Cluster × Treatment | 1.2363 | 2 | 0.5389 | ns |
| Cluster × Section | 41.5338 | 22 | 0.0071 | ** |
| Treatment × Section | 0.0971 | 11 | >0.9999 | ns |
| Cluster × Treatment × Section | 13.5132 | 22 | 0.9179 | ns |

**Supplementary Table 7. GLMM with section in omnibus results for microglial cluster proportions across 12 rostro-caudal hypothalamic sections in female embryos**

| Female Microglial Cluster Proportions (section in omnibus) | | |
|---|---|---|
| **Sidak-adjusted post hoc comparisons** | **Adjusted p value** | **Summary** |
| **Section 01 (females)** | | |
| Ameboid: Control vs. Cold Stress | >0.9999 | ns |
| Ramified: Control vs. Cold Stress | >0.9999 | ns |
| Rod-Like: Control vs. Cold Stress | 0.9158 | ns |
| **Section 02 (females)** | | |
| Ameboid: Control vs. Cold Stress | >0.9999 | ns |
| Ramified: Control vs. Cold Stress | >0.9999 | ns |
| Rod-Like: Control vs. Cold Stress | >0.9999 | ns |
| **Section 03 (females)** | | |
| Ameboid: Control vs. Cold Stress | 0.6369 | ns |
| Ramified: Control vs. Cold Stress | 0.5705 | ns |
| Rod-Like: Control vs. Cold Stress | >0.9999 | ns |
| **Section 04 (females)** | | |
| Ameboid: Control vs. Cold Stress | 0.9594 | ns |
| Ramified: Control vs. Cold Stress | 0.9958 | ns |
| Rod-Like: Control vs. Cold Stress | >0.9999 | ns |
| **Section 05 (females)** | | |
| Ameboid: Control vs. Cold Stress | >0.9999 | ns |
| Ramified: Control vs. Cold Stress | >0.9999 | ns |
| Rod-Like: Control vs. Cold Stress | >0.9999 | ns |
| **Section 06 (females)** | | |
| Ameboid: Control vs. Cold Stress | 0.9609 | ns |
| Ramified: Control vs. Cold Stress | >0.9999 | ns |
| Rod-Like: Control vs. Cold Stress | >0.9999 | ns |
| **Section 07 (females)** | | |
| Ameboid: Control vs. Cold Stress | 0.7788 | ns |
| Ramified: Control vs. Cold Stress | >0.9999 | ns |
| Rod-Like: Control vs. Cold Stress | 0.9999 | ns |
| **Section 08 (females)** | | |
| Ameboid: Control vs. Cold Stress | 0.9987 | ns |
| Ramified: Control vs. Cold Stress | >0.9999 | ns |
| Rod-Like: Control vs. Cold Stress | >0.9999 | ns |
| **Section 09 (females)** | | |
| Ameboid: Control vs. Cold Stress | >0.9999 | ns |
| Ramified: Control vs. Cold Stress | >0.9999 | ns |
| Rod-Like: Control vs. Cold Stress | >0.9999 | ns |
| **Section 10 (females)** | | |
| Ameboid: Control vs. Cold Stress | >0.9999 | ns |
| Ramified: Control vs. Cold Stress | >0.9999 | ns |

| | | |
|---|---|---|
| Rod-Like: Control vs. Cold Stress | >0.9999 | ns |
| **Section 11 (females)** | | |
| Ameboid: Control vs. Cold Stress | >0.9999 | ns |
| Ramified: Control vs. Cold Stress | >0.9999 | ns |
| Rod-Like: Control vs. Cold Stress | >0.9999 | ns |
| **Section 12 (females)** | | |
| Ameboid: Control vs. Cold Stress | >0.9999 | ns |
| Ramified: Control vs. Cold Stress | >0.9999 | ns |
| Rod-Like: Control vs. Cold Stress | >0.9999 | ns |

| **Type II Wald Chi-square tests** | **Chi-sq** | **Df** | **p value** | **Summary** |
|---|---|---|---|---|
| Cluster | 330.3624 | 2 | <0.0001 | **** |
| Treatment | 0.0279 | 1 | 0.8674 | ns |
| Section | 0.0803 | 11 | >0.9999 | ns |
| Cluster × Treatment | 13.9454 | 2 | 0.0009 | *** |
| Cluster × Section | 51.4863 | 22 | 0.0004 | *** |
| Treatment × Section | 0.0481 | 11 | >0.9999 | ns |
| Cluster × Treatment × Section | 24.8975 | 22 | 0.3020 | ns |

**Supplementary Table 8. Two-way ANOVA results for the number of microglial cells per mouse in the E15.5 PVN**

| **Number Microglial Cells in the PVN** | | |
|---|---|---|
| **Tukey's multiple comparisons test** | **Adjusted p value** | **Summary** |
| Male: Control vs. Male: Cold Stress | 0.8611 | ns |
| Male: Control vs. Female: Control | 0.3784 | ns |
| Male: Control vs. Female: Cold Stress | 0.7222 | ns |
| Male: Cold Stress vs. Female: Control | 0.1041 | ns |
| Male: Cold Stress vs. Female: Cold Stress | 0.2872 | ns |
| Female: Control vs. Female: Cold Stress | 0.9332 | ns |
| **Two-way ANOVA** | **p value** | **Summary** |
| Treatment | 0.3423 | ns |
| Sex | 0.0232 | * |
| Interaction | 0.8939 | ns |

**Supplementary Table 9. Two-way ANOVA results for the number of microglial cells per mouse in the E15.5 ARC**

| Number of Microglial Cells in the ARC | | |
|---|---|---|
| **Tukey's multiple comparisons test** | | |
| **Comparison** | **Adjusted p value** | **Summary** |
| Male: Control vs. Male: Cold Stress | 0.9877 | ns |
| Male: Control vs. Female: Control | 0.9544 | ns |
| Male: Control vs. Female: Cold Stress | 0.6894 | ns |
| Male: Cold Stress vs. Female: Control | 0.9975 | ns |
| Male: Cold Stress vs. Female: Cold Stress | 0.8621 | ns |
| Female: Control vs. Female: Cold Stress | 0.9335 | ns |
| **Two-way ANOVA** | **p value** | **Summary** |
| Treatment | 0.5238 | ns |
| Sex | 0.3701 | ns |
| Interaction | 0.8530 | ns |

**Supplementary Table 10. Two-way ANOVA results for microglia–AVP neuron touching events in the E15.5 control PVN**

| Microglia-AVP Neuron Touching Events in the PVN of Controls | | |
|---|---|---|
| **Tukey's multiple comparisons** | **Adjusted p value** | **Summary** |
| Ameboid: Male vs. Ameboid: Female | ns | 0.8958 |
| Ameboid: Male vs. Ramified: Male | ns | 0.5707 |
| Ameboid: Male vs. Ramified: Female | ns | 0.9996 |
| Ameboid: Male vs. Rodlike: Male | ns | 0.9734 |
| Ameboid: Male vs. Rodlike: Female | ns | 0.8318 |
| Ameboid: Female vs. Ramified: Male | ns | 0.1033 |
| Ameboid: Female vs. Ramified: Female | ns | 0.7532 |
| Ameboid: Female vs. Rodlike: Male | ns | 0.9996 |
| Ameboid: Female vs. Rodlike: Female | ns | >0.9999 |
| Ramified: Male vs. Ramified: Female | ns | 0.7532 |
| Ramified: Male vs. Rodlike: Male | ns | 0.1861 |
| Ramified: Male vs. Rodlike: Female | ns | 0.0751 |
| Ramified: Female vs. Rodlike: Male | ns | 0.8958 |
| Ramified: Female vs. Rodlike: Female | ns | 0.6643 |
| Rodlike: Male vs. Rodlike: Female | ns | 0.9974 |
| **Two-way ANOVA** | **p value** | **Summary** |
| Sex | 0.1094 | ns |
| Cluster | 0.0248 | * |
| Interaction | 0.8108 | ns |

**Supplementary Table 11. Two-way ANOVA results for microglia–AVP neuron wrapping events in the E15.5 control PVN**

| Microglia-AVP Neuron Wrapping Events in the PVN of Controls | | |
|---|---|---|
| **Tukey's multiple comparisons** | **Adjusted p value** | **Summary** |
| Ameboid: Male vs. Ameboid: Female | ns | 0.8958 |
| Ameboid: Male vs. Ramified: Male | ns | 0.5707 |
| Ameboid: Male vs. Ramified: Female | ns | 0.9996 |
| Ameboid: Male vs. Rodlike: Male | ns | 0.9734 |
| Ameboid: Male vs. Rodlike: Female | ns | 0.8318 |
| Ameboid: Female vs. Ramified: Male | ns | 0.1033 |
| Ameboid: Female vs. Ramified: Female | ns | 0.7532 |
| Ameboid: Female vs. Rodlike: Male | ns | 0.9996 |
| Ameboid: Female vs. Rodlike: Female | ns | >0.9999 |
| Ramified: Male vs. Ramified: Female | ns | 0.7532 |
| Ramified: Male vs. Rodlike: Male | ns | 0.1861 |
| Ramified: Male vs. Rodlike: Female | ns | 0.0751 |
| Ramified: Female vs. Rodlike: Male | ns | 0.8958 |
| Ramified: Female vs. Rodlike: Female | ns | 0.6643 |
| Rodlike: Male vs. Rodlike: Female | ns | 0.9974 |
| **Two-way ANOVA** | **p value** | **Summary** |
| Sex | 0.0154 | * |
| Cluster | 0.3709 | ns |
| Interaction | 0.5319 | ns |

**Supplementary Table 12. Two-way ANOVA results for microglia–AVP neuron touching events in the E15.5 male PVN**

| **Microglia-AVP Neuron Touching Events in the PVN of Males** | | |
|---|---|---|
| **Tukey’s multiple comparisons** | **Adjusted p value** | **Summary** |
| Ameboid: Cold Stress vs. Ameboid: Control | ns | 0.9995 |
| Ameboid: Cold Stress vs. Ramified: Cold Stress | * | 0.0349 |
| Ameboid: Cold Stress vs. Ramified: Control | ns | 0.8697 |
| Ameboid: Cold Stress vs. Rod-Like: Cold Stress | ns | >0.9999 |
| Ameboid: Cold Stress vs. Rod-Like: Control | ns | 0.9965 |
| Ameboid: Control vs. Ramified: Cold Stress | * | 0.0161 |
| Ameboid: Control vs. Ramified: Control | ns | 0.7039 |
| Ameboid: Control vs. Rod-Like: Cold Stress | ns | >0.9999 |
| Ameboid: Control vs. Rod-Like: Control | ns | >0.9999 |
| Ramified: Cold Stress vs. Ramified: Control | ns | 0.3252 |
| Ramified: Cold Stress vs. Rod-Like: Cold Stress | * | 0.0239 |
| Ramified: Cold Stress vs. Rod-Like: Control | * | 0.0108 |
| Ramified: Control vs. Rod-Like: Cold Stress | ns | 0.7939 |
| Ramified: Control vs. Rod-Like: Control | ns | 0.6058 |
| Rod-Like: Cold Stress vs. Rod-Like: Control | ns | 0.9995 |
| **Two-way ANOVA** | **p value** | **Summary** |
| Treatment | 0.1271 | ns |
| Cluster | 0.0020 | ** |
| Interaction | 0.3691 | ns |

**Supplementary Table 13. Two-way ANOVA results for microglia–AVP neuron wrapping events in the E15.5 male PVN**

| **Microglia-AVP Neuron Wrapping Events in the male PVN** | | |
|---|---|---|
| **Tukey's post hoc comparisons** | **Adjusted p value** | **Summary** |
| Ameboid: Cold Stress vs. Ameboid: Control | >0.9999 | ns |
| Ameboid: Cold Stress vs. Ramified: Cold Stress | 0.0001 | *** |
| Ameboid: Cold Stress vs. Ramified: Control | 0.9310 | ns |
| Ameboid: Cold Stress vs. Rod-Like: Cold Stress | >0.9999 | ns |
| Ameboid: Cold Stress vs. Rod-Like: Control | 0.9310 | ns |
| Ameboid: Control vs. Ramified: Cold Stress | <0.0001 | **** |
| Ameboid: Control vs. Ramified: Control | 0.8627 | ns |
| Ameboid: Control vs. Rod-Like: Cold Stress | >0.9999 | ns |
| Ameboid: Control vs. Rod-Like: Control | 0.9726 | ns |
| Ramified: Cold Stress vs. Ramified: Control | 0.0018 | ** |
| Ramified: Cold Stress vs. Rod-Like: Cold Stress | 0.0001 | *** |
| Ramified: Cold Stress vs. Rod-Like: Control | <0.0001 | **** |
| Ramified: Control vs. Rod-Like: Cold Stress | 0.9310 | ns |
| Ramified: Control vs. Rod-Like: Control | 0.4262 | ns |
| Rod-Like: Cold Stress vs. Rod-Like: Control | 0.9310 | ns |
| **Two-way ANOVA** | **p value** | **Summary** |
| Treatment | 0.0035 | ** |
| Cluster | <0.0001 | **** |
| Interaction | 0.0142 | * |

**Supplementary Table 14. Two-way ANOVA results for microglia–AVP neuron touching events in the E15.5 female PVN**

| **Microglia-AVP Neuron Touching Events in the female PVN** | | |
|---|---|---|
| **Tukey's post hoc comparisons** | **Adjusted p value** | **Summary** |
| Ameboid: Cold Stress vs. Ameboid: Control | 0.9998 | ns |
| Ameboid: Cold Stress vs. Ramified: Cold Stress | 0.0276 | * |
| Ameboid: Cold Stress vs. Ramified: Control | 0.5337 | ns |
| Ameboid: Cold Stress vs. Rod-Like: Cold Stress | >0.9999 | ns |
| Ameboid: Cold Stress vs. Rod-Like: Control | 0.9986 | ns |
| Ameboid: Control vs. Ramified: Cold Stress | 0.0511 | ns |
| Ameboid: Control vs. Ramified: Control | 0.6970 | ns |
| Ameboid: Control vs. Rod-Like: Cold Stress | 0.9998 | ns |
| Ameboid: Control vs. Rod-Like: Control | 0.9851 | ns |
| Ramified: Cold Stress vs. Ramified: Control | 0.6161 | ns |
| Ramified: Cold Stress vs. Rod-Like: Cold Stress | 0.0276 | * |
| Ramified: Cold Stress vs. Rod-Like: Control | 0.0104 | * |
| Ramified: Control vs. Rod-Like: Cold Stress | 0.5337 | ns |
| Ramified: Control vs. Rod-Like: Control | 0.3097 | ns |
| Rod-Like: Cold Stress vs. Rod-Like: Control | 0.9986 | ns |
| **Two-way ANOVA** | **p value** | **Summary** |
| Treatment | 0.3303 | ns |
| Cluster | 0.0010 | ** |
| Interaction | 0.4273 | ns |

**Supplementary Table 15. Two-way ANOVA results for microglia–AVP neuron wrapping events in the E15.5 female PVN**

| Microglia-AVP+ Neuron Wrapping Events in the Female PVN | | |
|---|---|---|
| **Tukey's post hoc comparisons** | **Adjusted p value** | **Summary** |
| Ameboid: Cold Stress vs. Ameboid: Control | >0.9999 | ns |
| Ameboid: Cold Stress vs. Ramified: Cold Stress | 0.4344 | ns |
| Ameboid: Cold Stress vs. Ramified: Control | >0.9999 | ns |
| Ameboid: Cold Stress vs. Rod-Like: Cold Stress | 0.9330 | ns |
| Ameboid: Cold Stress vs. Rod-Like: Control | >0.9999 | ns |
| Ameboid: Control vs. Ramified: Cold Stress | 0.3209 | ns |
| Ameboid: Control vs. Ramified: Control | >0.9999 | ns |
| Ameboid: Control vs. Rod-Like: Cold Stress | 0.9765 | ns |
| Ameboid: Control vs. Rod-Like: Control | >0.9999 | ns |
| Ramified: Cold Stress vs. Ramified: Control | 0.4344 | ns |
| Ramified: Cold Stress vs. Rod-Like: Cold Stress | 0.0820 | ns |
| Ramified: Cold Stress vs. Rod-Like: Control | 0.3755 | ns |
| Ramified: Control vs. Rod-Like: Cold Stress | 0.9330 | ns |
| Ramified: Control vs. Rod-Like: Control | >0.9999 | ns |
| Rod-Like: Cold Stress vs. Rod-Like: Control | 0.9585 | ns |
| **Two-way ANOVA** | **p value** | **Summary** |
| Treatment | 0.4754 | ns |
| Cluster | 0.1214 | ns |
| Interaction | 0.1713 | ns |

**Supplementary Table 16. Two-way ANOVA results for phagocytic microglia numbers by cluster in the E15.5 control PVN**

| **Microglial Phagocytosis in Controls** | | |
|---|---|---|
| **Tukey's multiple comparisons** | **Adjusted p value** | **Summary** |
| Ameboid: Male vs. Ameboid: Female | ns | >0.9999 |
| Ameboid: Male vs. Ramified: Male | ns | 0.1826 |
| Ameboid: Male vs. Ramified: Female | *** | 0.0002 |
| Ameboid: Male vs. Rodlike: Male | ns | 0.8293 |
| Ameboid: Male vs. Rodlike: Female | ns | 0.9838 |
| Ameboid: Female vs. Ramified: Male | ns | 0.1826 |
| Ameboid: Female vs. Ramified: Female | *** | 0.0002 |
| Ameboid: Female vs. Rodlike: Male | ns | 0.8293 |
| Ameboid: Female vs. Rodlike: Female | ns | 0.9838 |
| Ramified: Male vs. Ramified: Female | ns | 0.1081 |
| Ramified: Male vs. Rodlike: Male | * | 0.0123 |
| Ramified: Male vs. Rodlike: Female | * | 0.0450 |
| Ramified: Female vs. Rodlike: Male | **** | <0.0001 |
| Ramified: Female vs. Rodlike: Female | **** | <0.0001 |
| Rodlike: Male vs. Rodlike: Female | ns | 0.9941 |
| **Two-way ANOVA** | **p value** | **Summary** |
| Sex | 0.0726 | ns |
| Cluster | <0.0001 | **** |
| Interaction | 0.1504 | ns |

**Supplementary Table 17. Two-way ANOVA results for phagocytic microglia numbers by cluster in the E15.5 male PVN**

| **Microglial Phagocytosis in Males** | | |
|---|---|---|
| **Tukey's post hoc comparisons** | **Adjusted p value** | **Summary** |
| Ameboid: Cold Stress vs. Ameboid: Control | 0.9992 | ns |
| Ameboid: Cold Stress vs. Ramified: Cold Stress | 0.2944 | ns |
| Ameboid: Cold Stress vs. Ramified: Control | <0.0001 | **** |
| Ameboid: Cold Stress vs. Rod-Like: Cold Stress | 0.8891 | ns |
| Ameboid: Cold Stress vs. Rod-Like: Control | 0.8383 | ns |
| Ameboid: Control vs. Ramified: Cold Stress | 0.1567 | ns |
| Ameboid: Control vs. Ramified: Control | <0.0001 | **** |
| Ameboid: Control vs. Rod-Like: Cold Stress | 0.9788 | ns |
| Ameboid: Control vs. Rod-Like: Control | 0.9589 | ns |
| Ramified: Cold Stress vs. Ramified: Control | 0.0254 | * |
| Ramified: Cold Stress vs. Rod-Like: Cold Stress | 0.0337 | * |
| Ramified: Cold Stress vs. Rod-Like: Control | 0.0254 | * |
| Ramified: Control vs. Rod-Like: Cold Stress | <0.0001 | **** |
| Ramified: Control vs. Rod-Like: Control | <0.0001 | **** |
| Rod-Like: Cold Stress vs. Rod-Like: Control | >0.9999 | ns |
| **Two-way ANOVA** | **p value** | **Summary** |
| Treatment | 0.1097 | ns |
| Cluster | <0.0001 | **** |
| Interaction | 0.0235 | * |

**Supplementary Table 18. Two-way ANOVA results for phagocytic microglia numbers in the E15.5 female PVN**

| Microglial Phagocytosis in Females | | |
|---|---|---|
| **Tukey's post hoc comparisons** | **Adjusted p value** | **Summary** |
| Ameboid: Cold Stress vs. Ameboid: Control | 0.9884 | ns |
| Ameboid: Cold Stress vs. Ramified: Cold Stress | 0.0053 | ** |
| Ameboid: Cold Stress vs. Ramified: Control | 0.0420 | * |
| Ameboid: Cold Stress vs. Rod-Like: Cold Stress | 0.9950 | ns |
| Ameboid: Cold Stress vs. Rod-Like: Control | 0.6942 | ns |
| Ameboid: Control vs. Ramified: Cold Stress | 0.0257 | * |
| Ameboid: Control vs. Ramified: Control | 0.1580 | ns |
| Ameboid: Control vs. Rod-Like: Cold Stress | 0.8593 | ns |
| Ameboid: Control vs. Rod-Like: Control | 0.3251 | ns |
| Ramified: Cold Stress vs. Ramified: Control | 0.9592 | ns |
| Ramified: Cold Stress vs. Rod-Like: Cold Stress | 0.0013 | ** |
| Ramified: Cold Stress vs. Rod-Like: Control | 0.0001 | *** |
| Ramified: Control vs. Rod-Like: Cold Stress | 0.0119 | * |
| Ramified: Control vs. Rod-Like: Control | 0.0010 | ** |
| Rod-Like: Cold Stress vs. Rod-Like: Control | 0.9340 | ns |
| **Two-way ANOVA** | **p value** | **Summary** |
| Treatment | 0.5143 | ns |
| Cluster | <0.0001 | **** |
| Interaction | 0.4767 | ns |

**Supplementary Table 19. Two-way ANOVA results for microglial phagocytosis of bioparticles in the E15.5 hypothalamus**

| **Microglial Phagocytosis in the Hypothalamus** | | |
|---|---|---|
| **Tukey's post hoc comparisons** | **Adjusted p value** | **Summary** |
| Male: Control vs. Male: Cold Stress | 0.0379 | * |
| Male: Control vs. Female: Control | 0.6033 | ns |
| Male: Control vs. Female: Cold Stress | 0.4799 | ns |
| Male: Cold Stress vs. Female: Control | 0.3734 | ns |
| Male: Cold Stress vs. Female: Cold Stress | 0.5104 | ns |
| Female: Control vs. Female: Cold Stress | 0.9962 | ns |
| **Two-way ANOVA** | **p value** | **Summary** |
| Treatment | 0.0385 | * |
| Sex | 0.9610 | ns |
| Interaction | 0.0736 | ns |
