## Supplementary Figure 1-5 for "Fetal microglia show region-specific and morphology-dependent sex differences in their responsiveness to prenatal maternal stress"

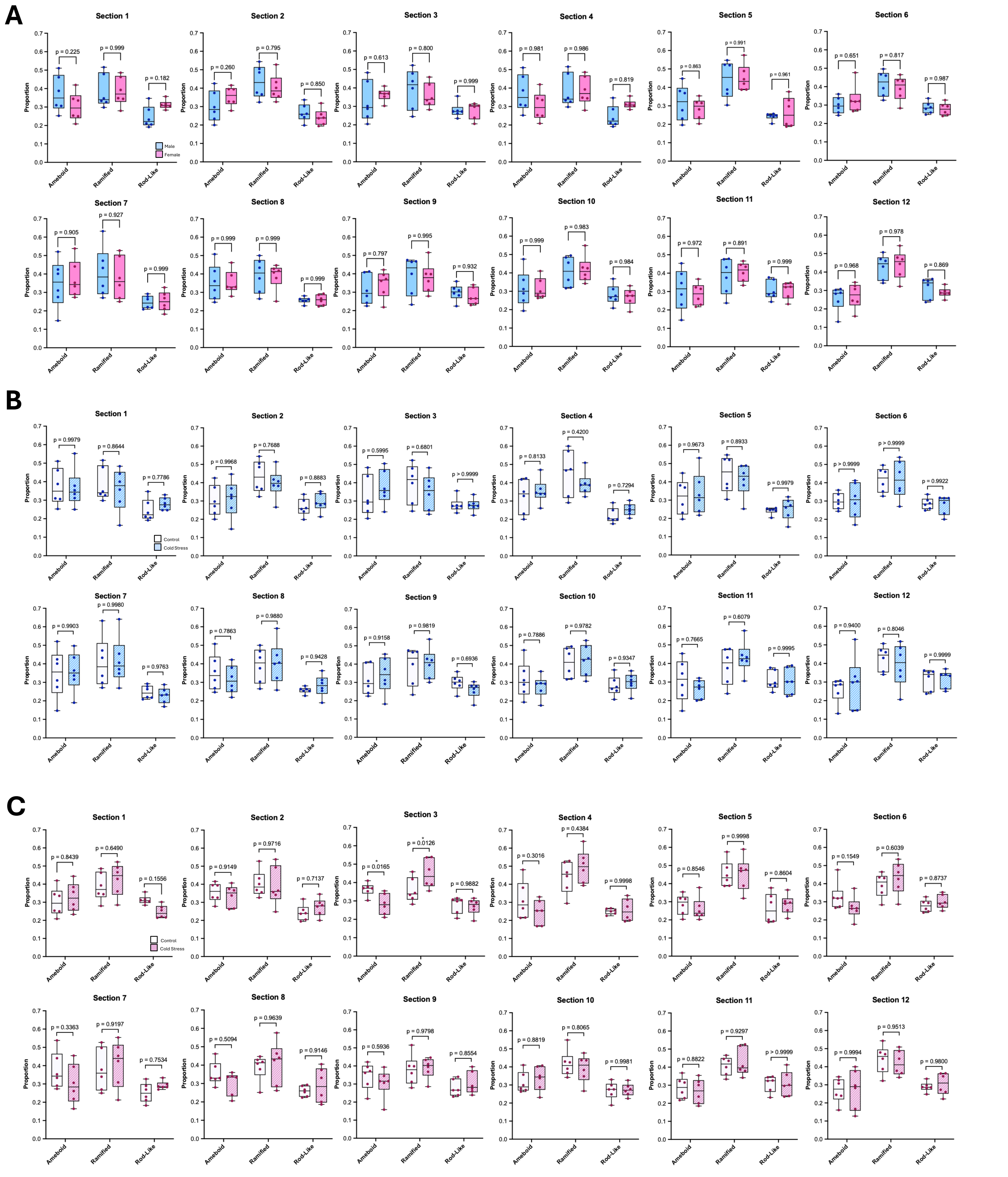
A
Section 1
p = 0.225
p = 0.999
p = 0.182
Male
Female
Section 2
p = 0.260
p = 0.795
p = 0.850
Section 3
p = 0.613
p = 0.800
p = 0.999
Section 4
p = 0.981
p = 0.986
p = 0.819
Section 5
p = 0.863
p = 0.991
p = 0.961
Section 6
p = 0.651
p = 0.817
p = 0.987
Section 7
p = 0.905
p = 0.927
p = 0.999
Section 8
p = 0.999
p = 0.999
p = 0.999
Section 9
p = 0.797
p = 0.995
p = 0.932
Section 10
p = 0.999
p = 0.983
p = 0.984
Section 11
p = 0.972
p = 0.891
p = 0.999
Section 12
p = 0.968
p = 0.978
p = 0.869
B
Section 1
p = 0.9979
p = 0.8644
p = 0.7786
Control
Cold Stress
Section 2
p = 0.9968
p = 0.7688
p = 0.8883
Section 3
p = 0.5995
p = 0.6801
p > 0.9999
Section 4
p = 0.8133
p = 0.4200
p = 0.7294
Section 5
p = 0.9673
p = 0.8933
p = 0.9979
Section 6
p > 0.9999
p > 0.9999
p = 0.9922
Section 7
p = 0.9903
p = 0.9980
p = 0.9763
Section 8
p = 0.7863
p = 0.9880
p = 0.9428
Section 9
p = 0.9158
p = 0.9819
p = 0.6936
Section 10
p = 0.7886
p = 0.9782
p = 0.9347
Section 11
p = 0.7665
p = 0.6079
p = 0.9995
Section 12
p = 0.9400
p = 0.8046
p = 0.9999
C
Section 1
p = 0.8439
p = 0.6490
p = 0.1556
Control
Cold Stress
Section 2
p = 0.9149
p = 0.9716
p = 0.7137
Section 3
*
p = 0.0165
*
p = 0.0126
p = 0.9882
Section 4
p = 0.3016
p = 0.4384
p = 0.9998
Section 5
p = 0.8546
p = 0.9998
p = 0.8604
Section 6
p = 0.1549
p = 0.6039
p = 0.8737
Section 7
p = 0.3363
p = 0.9197
p = 0.7534
Section 8
p = 0.5094
p = 0.9639
p = 0.9146
Section 9
p = 0.5936
p = 0.9798
p = 0.8554
Section 10
p = 0.8819
p = 0.8065
p = 0.9981
Section 11
p = 0.8822
p = 0.9297
p > 0.9999
Section 12
p = 0.9994
p = 0.9513
p = 0.9800
Proportion
Ameboid
Ramified
Rod-Like

**Supplementary Figure 1. Microglial cluster prevalence across the rostral–caudal axis of the E15.5 hypothalamus.** (A) Proportion of each microglial cluster (ameboid vs. ramified vs. rod-like) across the rostral–caudal axis of the E15.5 control male and female hypothalamus. (B) Proportion of each microglial cluster across the rostral–caudal axis of the E15.5 control and cold stress male hypothalamus. (C) Proportion of each microglial cluster across the rostral–caudal axis of the E15.5 control and cold stress female hypothalamus. Microglial cluster proportions shifted in section 3 (cluster, $p<0.0001$; treatment, $p=0.9761$; cluster×treatment interaction, $p=0.0003$), with a significant reduction in ameboid microglia ($p=0.0165$) and a significant increase in ramified microglia ($p=0.0126$) in cold stress females. (A–C) Data are represented as box-and-whisker plots showing the median (center line), interquartile range (box), and minimum–maximum values (whiskers) and were analyzed by a linear mixed-effects model with Sidak-adjusted post hoc comparisons for each section. Each dot represents an individual embryo (n=6 embryos), which were collected from 3 independent litters.

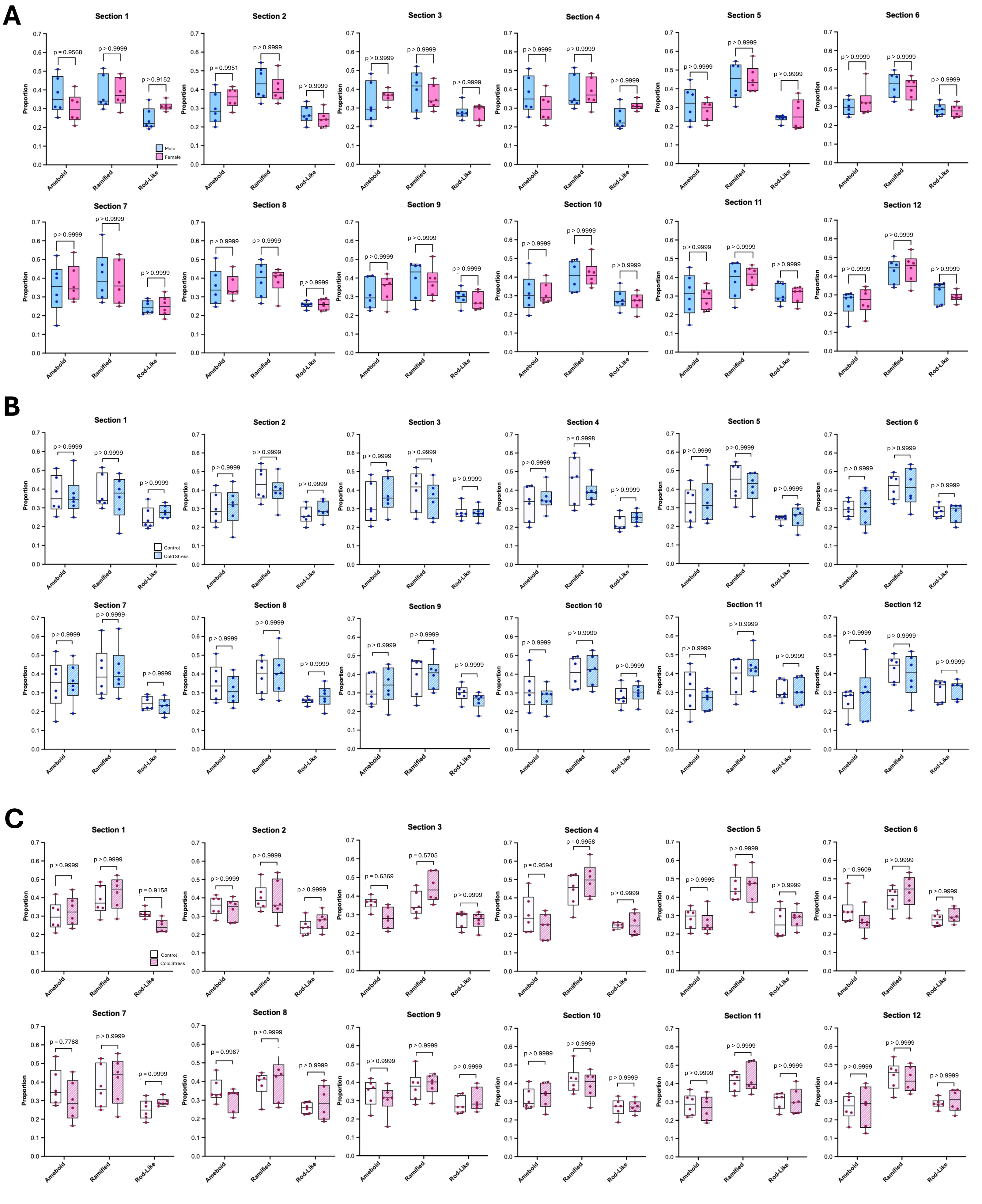
A
Male
Female
Section 1
p = 0.9568
p > 0.9999
p = 0.9152
Section 2
p = 0.9951
p > 0.9999
p > 0.9999
Section 3
p > 0.9999
p > 0.9999
p > 0.9999
Section 4
p > 0.9999
p > 0.9999
p > 0.9999
Section 5
p > 0.9999
p > 0.9999
p > 0.9999
Section 6
p > 0.9999
p > 0.9999
p > 0.9999
Section 7
p > 0.9999
p > 0.9999
p > 0.9999
Section 8
p > 0.9999
p > 0.9999
p > 0.9999
Section 9
p > 0.9999
p > 0.9999
p > 0.9999
Section 10
p > 0.9999
p > 0.9999
p > 0.9999
Section 11
p > 0.9999
p > 0.9999
p > 0.9999
Section 12
p > 0.9999
p > 0.9999
p > 0.9999
B
Control
Cold Stress
Section 1
p > 0.9999
p > 0.9999
p > 0.9999
Section 2
p > 0.9999
p > 0.9999
p > 0.9999
Section 3
p > 0.9999
p > 0.9999
p > 0.9999
Section 4
p > 0.9999
p = 0.9998
p > 0.9999
Section 5
p > 0.9999
p > 0.9999
p > 0.9999
Section 6
p > 0.9999
p > 0.9999
p > 0.9999
Section 7
p > 0.9999
p > 0.9999
p > 0.9999
Section 8
p > 0.9999
p > 0.9999
p > 0.9999
Section 9
p > 0.9999
p > 0.9999
p > 0.9999
Section 10
p > 0.9999
p > 0.9999
p > 0.9999
Section 11
p > 0.9999
p > 0.9999
p > 0.9999
Section 12
p > 0.9999
p > 0.9999
p > 0.9999
C
Control
Cold Stress
Section 1
p > 0.9999
p > 0.9999
p = 0.9158
Section 2
p > 0.9999
p > 0.9999
p > 0.9999
Section 3
p = 0.6369
p = 0.5705
p > 0.9999
Section 4
p = 0.9594
p = 0.9958
p > 0.9999
Section 5
p > 0.9999
p > 0.9999
p > 0.9999
Section 6
p = 0.9609
p > 0.9999
p > 0.9999
Section 7
p = 0.7788
p > 0.9999
p > 0.9999
Section 8
p = 0.9987
p > 0.9999
p > 0.9999
Section 9
p > 0.9999
p > 0.9999
p > 0.9999
Section 10
p > 0.9999
p > 0.9999
p > 0.9999
Section 11
p > 0.9999
p > 0.9999
p > 0.9999
Section 12
p > 0.9999
p > 0.9999
p > 0.9999
Proportion
0.0
0.1
0.2
0.3
0.4
0.5
0.6
0.7
Ameboid
Ramified
Rod-Like

**Supplementary Figure 2. Microglial cluster prevalence across the rostral–caudal axis of the E15.5 hypothalamus with hypothalamic section in the omnibus.** (A) Proportion of each microglial cluster (ameboid vs. ramified vs. rod-like) across the rostral–caudal axis of the E15.5 control male and female hypothalamus. (B) Proportion of each microglial cluster across the rostral–caudal axis of the E15.5 control and cold stress male hypothalamus. (C) Proportion of each microglial cluster across the rostral–caudal axis of the E15.5 control and cold stress female hypothalamus. (A–C) Data are represented as box-and-whisker plots showing the median (center line), interquartile range (box), and minimum–maximum values (whiskers) and were analyzed by a linear mixed-effects model with Sidak-adjusted post hoc comparisons for each cluster and section. Each dot represents an individual embryo (n=6 embryos), which were collected from 3 independent litters.

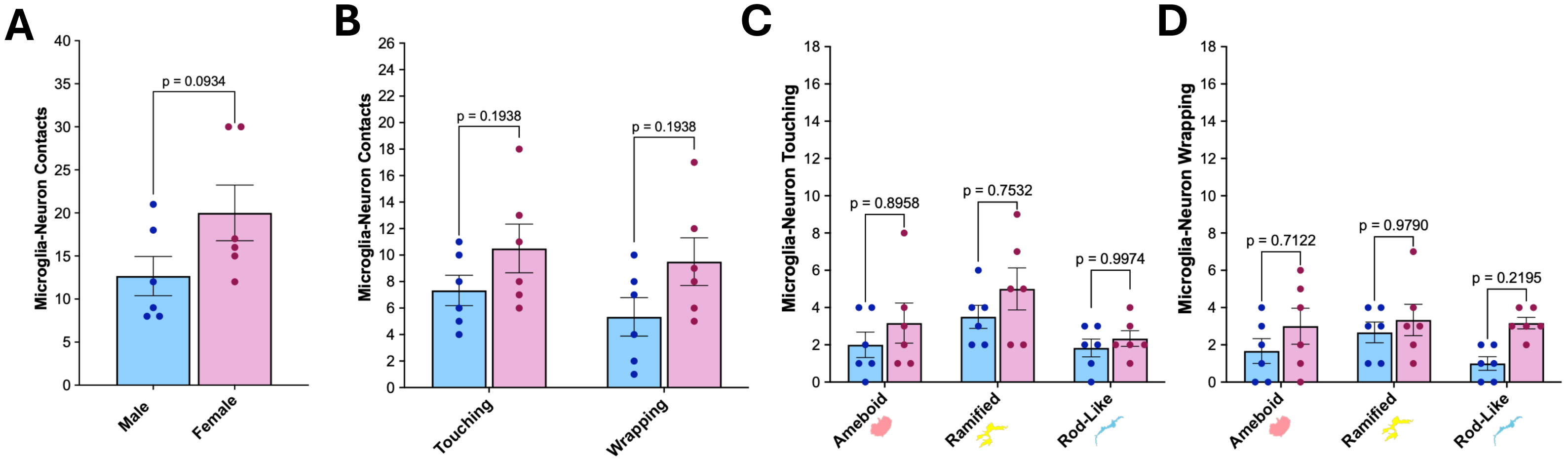
A
Microglia-Neuron Contacts
p = 0.0934
Male
Female
B
Microglia-Neuron Contacts
p = 0.1938
p = 0.1938
Touching
Wrapping
C
Microglia-Neuron Touching
p = 0.8958
p = 0.7532
p = 0.9974
Ameboid
Ramified
Rod-Like
D
Microglia-Neuron Wrapping
p = 0.7122
p = 0.9790
p = 0.2195
Ameboid
Ramified
Rod-Like

**Supplementary Figure 3. Microglia–AVP neuron contacts in the E15.5 control male and female PVN.** (A–B) Total microglia–AVP neuron contacts (A) and contacts by event type (touching vs. wrapping) (B) in the E15.5 control male and female PVN. (C) Microglia–AVP neuron touching events by cluster (ameboid vs. ramified vs. rod-like) in the E15.5 control male and female PVN (cluster, $p=0.0248$; sex, $p=0.1094$; cluster×sex interaction, $p=0.8108$). (D) Microglia–AVP neuron wrapping events by cluster in the E15.5 control male and female PVN (cluster, $p=0.3709$; sex, $p=0.0154$; cluster×sex interaction, $p=0.5319$). (A–D) Data are represented as means ± SEM and were analyzed using a Student's t-test (A, B) or a two-way ANOVA with Tukey-adjusted post hoc comparisons (C, D). Each dot represents an individual embryo ($n=6$ embryos), which were collected from 3 independent litters.

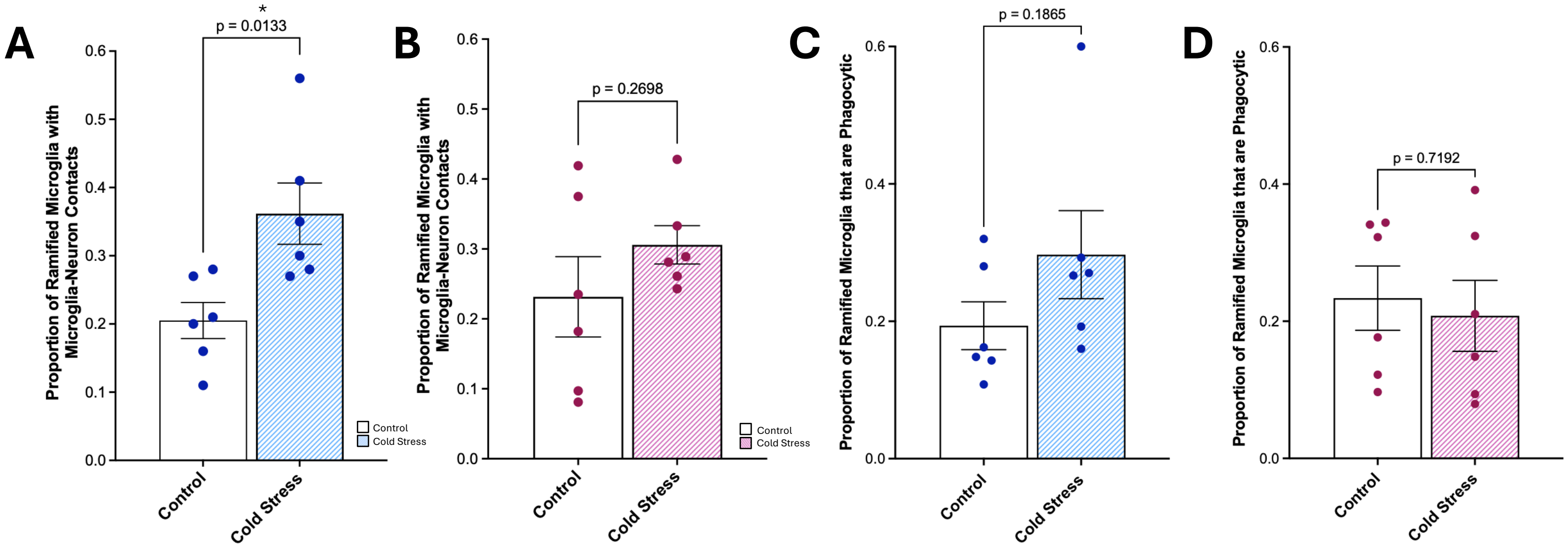

A
B
C
D
Proportion of Ramified Microglia with Microglia-Neuron Contacts
Proportion of Ramified Microglia that are Phagocytic
p = 0.0133
*
p = 0.2698
p = 0.1865
p = 0.7192
Control
Cold Stress

**Supplementary Figure 4. Proportion of ramified microglia contacting AVP neurons or engaging in phagocytosis in the E15.5 PVN.** (A, B) Proportion of ramified microglia contacting AVP neurons in the E15.5 control and cold stress male (A) and female (B) PVN. Cold stress males have significantly more ramified microglia–AVP neuron contacts compared to control males ($p=0.0133$). (C, D) Proportion of ramified microglia that are phagocytic in the E15.5 control and cold stress male (C) and female (D) PVN. (A–D) Data are represented as means ± SEM and were analyzed using a Student's t-test. Each dot represents an individual embryo ($n=6$ embryos), which were collected from 3 independent litters.

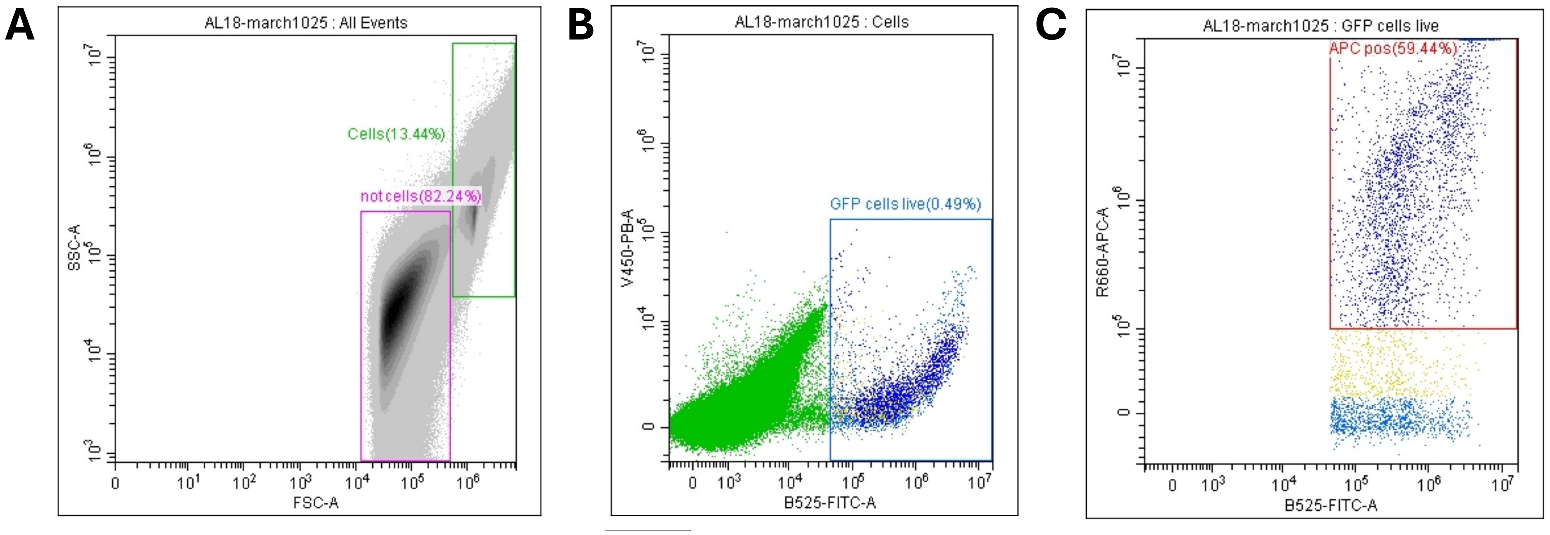
A
AL18-march1025 : All Events
Cells(13.44%)
not cells(82.24%)
SSC-A
FSC-A
B
AL18-march1025 : Cells
GFP cells live(0.49%)
V450-PB-A
B525-FITC-A
C
AL18-march1025 : GFP cells live
APC pos(59.44%)
R660-APC-A
B525-FITC-A

**Supplementary Figure 5. Flow cytometry gating strategy to quantify the proportion of microglia containing engulfed bioparticles in the E15.5 hypothalamus.** (A) Exclusion of debris, and identification of cells. (B) Gating of GFP+ microglia, within the cell population. (C) Gating of APC+ (bioparticle-containing) microglia, within the GFP+ population.
